# Incorporating photosynthetic acclimation improves stomatal optimisation models

**DOI:** 10.1101/2023.07.07.548137

**Authors:** Victor Flo, Jaideep Joshi, Manon Sabot, David Sandoval, Iain Colin Prentice

## Abstract

Stomatal opening in plant leaves is regulated through a balance of carbon and water exchange under different environmental conditions. Accurate estimation of stomatal regulation is crucial for understanding how plants respond to changing environmental conditions, particularly under climate change. A new generation of optimality-based modelling schemes determines instantaneous stomatal responses from a balance of trade-offs between carbon gains and hydraulic costs, but most such schemes do not account for biochemical acclimation in response to drought. Here, we compare the performance of seven instantaneous stomatal optimisation models with and without accounting for photosynthetic acclimation. Using experimental data from 38 plant species, we found that accounting for photosynthetic acclimation improves the prediction of carbon assimilation in a majority of the tested models. Non-stomatal mechanisms contributed significantly to the reduction of photosynthesis under drought conditions in all tested models. Drought effects on photosynthesis could not accurately be explained by the hydraulic impairment functions embedded in the stomatal models alone, indicating that photosynthetic acclimation must be considered to improve estimates of carbon assimilation during drought.

**Summary Statement:** Accounting for photosynthetic acclimation improves the predictions of carbon assimilation in all the stomatal optimization models evaluated. The influence of drought on photosynthesis cannot be fully explained by the hydraulic impairment function of the stomatal models alone.

## Introduction

Accurate estimation of net photosynthetic rates (A_net_) in terrestrial ecosystems is crucial for understanding and predicting vegetation dynamics, global carbon budgets, and climate change trends (Arora et al., 2020; Fisher & Koven, 2020; Green et al., 2019; Humphrey et al., 2021; Lombardozzi et al., 2015; Mercado et al., 2018). However, photosynthesis modules within current Ecosystem and Earth System models often lack accuracy and provide biased estimates of carbon assimilation (Prentice et al., 2015; Rogers et al., 2017; Seiler et al., 2022). This is explained, among other factors, by the inability of large-scale models to adequately account for the effects of drought on the regulation of stomatal gas exchange and leaf photosynthetic capacity (Zhou et al., 2013). Novel hydraulically explicit stomatal optimization models that mechanistically incorporate drought impacts have shown promise in improving the accuracy of instantaneous gas exchange predictions, and therefore A_net_, in terrestrial models (Eller et al., 2020; Sabot et al., 2020), while reducing their reliance on poorly defined empirical parameters. Previous studies have thoroughly evaluated the performance of a variety of such models (Sabot et al., 2022b; Wang et al., 2020), and have equivocally advocated the integration of non-stomatal limitations alongside stomatal limitations to photosynthesis, particularly, drought impacts on photosynthetic capacity.

Drought is a major abiotic stressor that affects plant growth and productivity (Ahlström et al., 2015; Kannenberg, Schwalm & Anderegg, 2020; Ruehr et al., 2019; Stocker et al., 2019). Under soil drought conditions or significantly increased atmospheric vapour pressure deficit (VPD), plants typically decrease their stomatal conductance (g_s_) to reduce water use and protect their hydraulic system from embolism. However, a decrease in g_s_ leads to a reduction in CO_2_ diffusion within the leaves, which in turn results in a decrease in A_net_. Therefore, the regulation of g_s_ entails a trade-off between uptake of CO_2_ for photosynthesis and managing hydraulic risks. As A_net_ declines during drought, maximum leaf photosynthetic capacity also goes down to reduce metabolic costs, such as those associated with ribulose-1,5-bisphosphate (RuBP) turnover and ATP synthesis (Flexas & Medrano, 2002; Martin-StPaul et al., 2012; Peguero-Pina et al., 2009; Salmon et al., 2020; Zhou et al., 2013). This could be an adaptation to minimise the investment of resources in unprofitable processes and instead diverting them towards growth and development to maximize fitness (Bloom, 1986, Anon, 2005). Downregulation of photosynthetic capacity has also been proposed to be associated with photochemical inhibition by elevated sugar concentrations in mesophyll cells (Hölttä et al., 2017), as well as increased abscisic acid accumulation (ABA) and electrolyte concentrations, inhibiting stromal enzymes (Kaiser, 1987). Although the specific proximate mechanisms governing drought-driven photosynthetic downregulation are not fully understood, it is plausible that its ultimate cause is the cost associated with maintaining photosynthetic capacity. Therefore, exploring its implications for stomatal control provides a basis for improving the accuracy of photosynthetic predictions (Sabot et al., 2022b; Smith et al., 2019; Yang et al., 2019).

Stomatal optimisation models rest on an optimal balance of the trade-offs between the carbon gained by photosynthesis and the risks associated with water lost to transpiration (Cowan & Farquhar, 1977). Inspired by economic thinking, such trade-off optimisation is typically expressed as a carbon profit equation (i.e., carbon gain minus carbon cost), where water loss is substituted for an equivalent carbon-based ‘penalty’ cost. Thus, plants may take the opportunity for higher carbon gain by allowing more transpiration when water is cheap to use and forego carbon gain when it is costly. A major appeal of optimality-based stomatal models is that, unlike their empirical counterparts (e.g., Ball, Woodrow & Berry, 1987; Leuning, 1995, 1990), they avoid prescribing the sensitivities of g_s_ to the environment and, instead, let them emerge from the optimality criterion (Medlyn et al., 2011; Prentice et al., 2014; Sperry et al., 2017). The added value in using optimisation models is robustness, i.e., the ability to predict stomatal behaviour under new environmental conditions outside of those used for model parameterization. The challenge however, is in formulating the optimality criterion and transforming water loss into a penalty cost, owing to a variety of ways in which transpiration and ensuing soil water depletion can impede physiological function (Wang et al., 2020). Following Cowan and Farquhar (1977), stomatal optimisation models originally considered a parametric marginal carbon cost of water use (Katul, Palmroth & Oren, 2009; Konrad, Roth-Nebelsick & Grein, 2008; Mäkelä, 1996; Manzoni et al., 2011; Medlyn et al., 2011). Yet, it was never specified how these costs should vary with the environment, time, and species (Manzoni et al., 2013, 2011; Schymanski et al., 2007; Wong, Cowan & Farquhar, 1985). To address this issue, subsequent stomatal optimization models have conceptualised the cost of water use as an impairment to hydraulic function arising from the cavitation of xylem vascular tissues (Anderegg et al., 2018; Buckley, Frehner & Bailey, 2023; Eller et al., 2020; Joshi et al., 2022; Lu et al., 2020; Sperry et al., 2017; Wang et al., 2020; Wolf, Anderegg & Pacala, 2016).

Motivated by calls to include non-stomatal limitations to photosynthesis in terrestrial models (Sun et al., 2014; Zhou et al., 2013), some studies have attempted to account for non-stomatal limitations within stomatal optimization criteria (Dewar et al., 2018; Hölttä et al., 2017). These studies attempted to capture gas exchange responses to variation in soil water availability by maximizing photosynthesis under a prescribed linear reduction in instantaneous photosynthetic capacity or mesophyll conductance with decreasing xylem pressure (Dewar et al., 2018) or decreasing turgor pressure (Hölttä et al., 2017). However, this assumption of linear downregulation is little supported by evidence (Wang et al., 2020). Other studies (Drake, 2017; Egea, Verhoef & Vidale, 2011; Keenan et al., 2009; Knauer et al., 2019; Yang et al., 2019; Zhou et al., 2014, 2013) have relied on calibrated functions for prescribing the reduction in maximum photosynthetic capacity or of the maximum mesophyll conductance. By contrast with the aforementioned approaches, Joshi *et al*. (2022) suggested to add metabolic impairment to hydraulic impairment within the stomatal optimization procedure, by accounting for a marginal maintenance cost of photosynthetic capacity (α). How α changes across space and through time (e.g., with environmental growing conditions, with changes in other functional traits) is unclear, but metabolic impairments are usually correctly represented in models when they are allowed to vary over timescales of a few days (Caldararu et al., 2020; Haverd et al., 2018; Jiang et al., 2020; Sabot et al., 2022a), whereas stomatal regulation is presumed instantaneous.

In this study, we investigated the effects of including an optimality-based representation of photosynthetic acclimation within six existing models and a new variant of one of them. To that end, we extended their optimality criteria to consider a marginal maintenance cost of photosynthetic capacity α, as proposed in Joshi *et al*. (2022). We tested the performance of these modified models against observational data stemming from dry-down experiments on >35 C_3_ species. Our main objective was to assess the ability of the extended models to predict A_net_ and g_s_, compared to the original formulations (Objective 1). We also analysed the extent to which each model’s estimates of A_net_ and g_s_ could be explained by hydraulic impairment vs. non-stomatal limitations manifested through the regulation of photosynthetic capacity (Objective 2). Finally, we explored the relationship between the cost parameter α, hydraulic parameters, (sub)species-specific functional traits, and their environmental growth conditions (Objective 3). We expected that accounting for photosynthetic acclimation will improve A_net_ estimates for all models. Based on previous findings by Wang *et al*. (2020) and Sabot *et al*. (2022b), we also expected that hydraulic impairment will affect assimilation to a greater extent than non-stomatal limitations during soil dry-down, albeit to varying degrees among models. Finally, we hypothesized that species characterized by high hydraulic vulnerability (e.g., high P_50_) or high specific leaf areas would have higher photosynthetic maintenance costs, and that “fast growing but drought sensitive” species would also have higher metabolic sensitivity to drought.

## Methods

### Stomatal optimization models

Hydraulics-enabled stomatal optimization models couple water transport and carbon uptake through g_s_ (see Notes 1 in the Supplementary Material). In these steady-state models, g_s_ links to leaf water potential (ψ_l_) through a mass-balance of water, i.e., root-to-leaf water supply equals transpiration demand (Eq. S5), whilst controlling the rate at which CO_2_ diffuses within the leaf, and thus, A_net_ (Eq. S6). A given model’s optimality criterion works to adjust the instantaneous g_s_ (consequently, ψ_l_) to maximize profit, i.e., A_net_ minus a carbon-equivalent hydraulic cost Θ (Eq. 1). Stomatal optimization models can be algebraically rearranged such that they differ only in the mathematical form of the Θ function (see Sabot et al., 2022b and Wang et al., 2020 for comprehensive reviews).

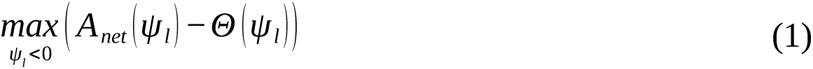

Adopting the nomenclature used by Sabot et al. (2022b), the stomatal optimisation models considered here are: PMAX (Sperry et al., 2017), PMAX2 (Wang et al., 2020), PMAX3 (this study), SOX (Eller et al., 2018), SOX2 (Buckley et al. 2023), CGAIN (Lu et al., 2020), and PHYDRO (Joshi et al., 2022). Notice that in this study we consider differences in water potential as equivalent to differences in water pressure (but see Wang & Frankenberg, 2022 for an appraisal of this assumption).

The PMAX model was proposed by Sperry et al. (2017) with the aim to express the stomatal cost of water loss based on plant hydraulic theory at a minimal parameter expense. The PMAX model balances a carbon gain function, where A_net_ is normalised by its maximum potential value (A_max_) specific to the instantaneous environmental conditions, with a hydraulic risk function, also normalised, varying with ψ_l_ and soil water potential (ψ_s_). A unique pairing of g_s_ and ψ_l_ hence maximises the difference between the normalised assimilation and risk functions. This model can also be expressed as in Equation 1 by multiplying both the gain function and hydraulic risk function by Amax, in which case Θ becomes

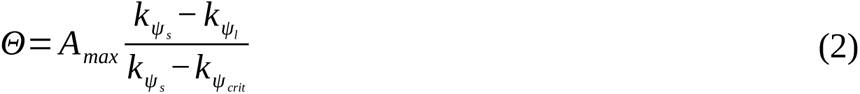

where *k*ψ is the xylem hydraulic conductance at a given water potential (ψ) calculated using Eq. S3 (Notes 1 in Supplementary Material), with the double-subscripts *s*, *l,* and *crit* representing soil, leaf, and critical water potentials, respectively. The hydraulic conductance at the critical water potential (*k*ψ*crit*) represents the point at which the plant pays the maximum cost of canopy desiccation (Sperry et al. 2017). Here, *k*ψ*crit* is assumed to be zero for simplicity and generalisation across species.

The PMAX2 model was developed by Wang et al. (2020) in order to meet a specific set of mathematical and biological criteria for a biologically sensible Θ. The hydraulic impairment function of this model is based on penalising the instantaneous transpiration (*E*(ψl)) by its proximity to *Ecrit,* i.e., *E*(ψl) at the critical leaf water potential at which we assume hydraulic conductance is null (ψl → –∞). *E*(ψl) is weighted by Anet in proportion to *Ecrit* such that

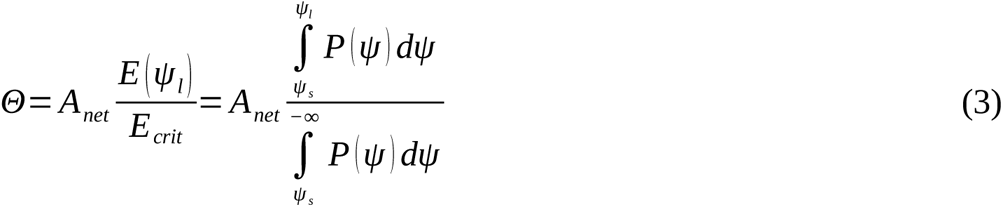

Here, the hydraulic impairment function is equivalent to the ratio _of_ two integrals of the hydraulic vulnerability curve function (P(ψ)). The integral in the numerator ranges from ψ_s_ to ψ_l_, and the improper integral in the denominator ranges from ψ_s_ to minus infinity (at *k*_ψ*crit*_). P(ψ) is calculated using Eq. S4.

The PMAX3 model proposed in this study is a biologically more consistent variant of the PMAX2 model. The model, instead of maximising the differential between A_net_ and the cost of hydraulic impairment, minimises the opportunity cost of carbon and the hydraulic cost. The model uses a similar hydraulic impairment function as PMAX2. Expressed as in eq. 1,

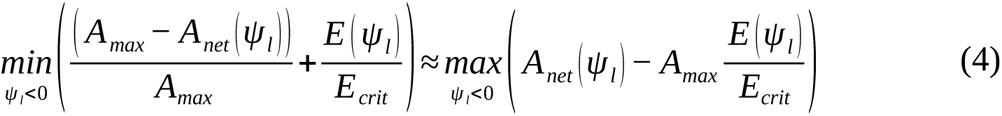

The CGAIN stomatal optimization model (Lu et al., 2020) was motivated by the fact that gas exchange may be regulated differently depending on dry-down length and dry-down interval time.

The model assumes a carbon cost (ϖ; μmol m^-2^ s^-1^) to offset carbon gain, with the former representing an investment into recovering the hydraulic conductance lost to embolized xylem (Sabot et al., 2022b). While the CGAIN model can account for the lagged effects of xylem recovery, as in Sabot *et al*. (2022b), we assume no lagged costs of xylem recovery. This simplified version of CGAIN is expressed as

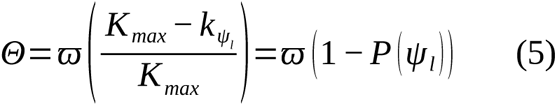

where K_max_ is the maximum whole-plant xylem hydraulic conductance between the roots to the leaves.

The PHYDRO model (Joshi et al. 2022) took a phenomenological approach that relies on an impairment function proportional to the square of the differential between the soil and leaf water potential (Δψ^2^)

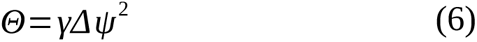

where γ (μmol m^-2^ s^-1^ MPa^-2^) is a calibrated empirical parameter which determines species-specific hydraulic sensitivity and is potentially related to the isohydricity of a species. This model requires the use of whole-plant vulnerability curve, including the outside-xylem hydraulic pathways in fine roots and leaves.

The SOX stomatal model (Eller et al. 2018) drew on the PMAX model but opted to directly impair A_net_ by the ratio of the hydraulic vulnerability curve evaluated at ψ_l_ to that of the vulnerability curve evaluated at ψ_s_

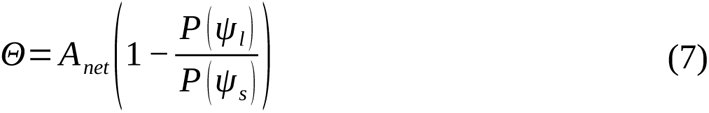

Finally, the SOX2 model proposed by Buckley et al. (2023) reworked the SOX model to account for the leaf physiological kinetic factors that affect leaf water potential dynamics. In this model the parameter ψ_50_ (i.e. the water potential at a 50% loss of xylem hydraulic conductance) is tuned by a species-specific stomatal behaviour parameter, δ (dimensionless). The parameter *b* (dimensionless) is the steepness of the vulnerability curve

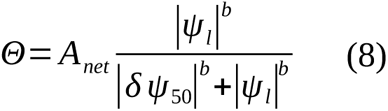

### Acclimation of the photosynthetic capacity

To calculate A_net_, maximum photosynthetic capacity must be known. Following the photosynthetic model of Farquhar *et al*. (1980), A_net_ is determined by the lesser of two biochemical assimilation rates (Aj and Ac) that are functions of the available irradiation (I_abs_), leaf intercellular CO_2_ concentration (C_i_), and temperature (Eq. S7 and Eq. S8), minus a day leaf respiration rate. The two rates Aj and Ac depend on the maximum rate of electron transport (J_max_) and maximum rate of carboxylation capacity (V_cmax_), respectively, with both J_max_ and V_cmax_ differing in their sensitivities to temperature. The parameter values that set J_max_ and V_cmax_ are typically standardized to 25°C (J_max,25_ and V_cmax,25_; Eqs. S14 ad S15) and taken to be species, ecosystem, or functional-type averages, without accounting for variation in their standardized magnitudes (i.e., the ratio of J_max,25_ to V_cmax,25_ is a fixed value). However, J_max_ and V_cmax_ have been observed to co-vary in the medium to long term, leading to a coordination among the two photosynthetic rates (Chen et al., 1993).

Assuming coordination, we can establish a single photosynthetic capacity cost, based on only one of the two limiting biochemical photosynthetic rates. Specifically, we let both J_max_ and V_cmax_ acclimate by extending the stomatal optimisation criterion (Eq. 1) to account for the cost of maintaining electron-transport capacity (Joshi et al. 2022).

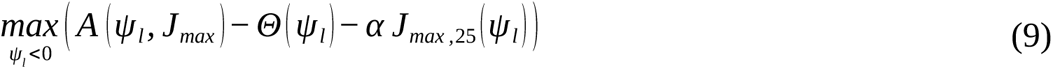

where α is a marginal maintenance cost per unit of electron-transport capacity under standard conditions (i.e. 25 °C), and αJ_max,25_ represents the total carbon cost required to maintain photosynthetic capacity. Here, unlike in Joshi et al. (2022), we optimise J_max,25_ instead of J_max_, to limit the effects of temperature on α.

The optimisation problem presented in Eq. 9 allows us to estimate the values of ψ_l_ and J_max,25_ that maximise profit, similar to how Eq. 1 allows us to regulate ψ_l_ to maximise A_net_. The optimisation of J_max,25_ is performed instantaneously, but using the average environmental conditions over the past seven days to capture the weekly acclimation timescale. We obtain the corresponding coordinated V_cmax,25_ via the photosynthetic coordination hypothesis (Eq. S10 – Eq. S13). The optimised acclimated parameter values of J_max,25_ and V_cmax,25_ are then adjusted to the instantaneous temperature using an Arrhenius function (Eqs. S14 and S15), and fed back as fixed parameters into the biochemical photosynthetic model for the calculation of instantaneous g_s_, ψ_s_, and A_net_ via Eq. 1.

In our approach, photosynthetic capacity is not only acclimated to the prevailing atmospheric conditions of the previous seven-day period, but also to ψ_s_ through the averaged hydraulic cost or impairment factor. However, as soil moisture conditions were not available for all seven day periods, we assumed that ψ_s_ decreased linearly between the first day of the experiment and the last day. We also assumed that the environmental conditions were constant until the start of the experiment.

### Experimental data

We used published data from a series of dry-down experiments to evaluate the performance of the seven stomatal optimization models with or without accounting for the photosynthetic acclimation. Some of these experimental data were originally compiled from the published literature by Zhou et al. (2013). We complemented them with others found through a similar literature search on Google Scholar – in September 2022, we used the keywords ‘“stomatal conductance” and “photosynthesis” and “soil water potential” and “dry-down” or “drought” or “experiment”’, yielding 490 references.

Data were subsequently requested from the authors, collected from open databases, or digitised from published figures using GetData Graph Digitiser version 2.26.0.20. The data were filtered to ensure that the experiments fulfilled the following criteria: 1) individuals of the species were subjected to a controlled drought, 2) the growth conditions were recorded during the study period, and 3) simultaneous measurements of g_s_, A_net_, and soil water potential or pre-dawn leaf water potential were made. For one of the experiments (Salmon et al., 2020), we also included the values of the controls where the individuals were not exposed to drought (i.e., the soil water potential was above –1 MPa) to expand data limitations at high soil-water potentials. After filtering, data from 13 studies remained, for a total of 38 species and sub-species representing diverse plant functional types (Table 1, Fig. S1-S2).

**Table 1.** Species and sub-species included in this study, as well as their hydraulic characteristics. ψ_50_ is the water potential at a 50% loss of xylem hydraulic conductivity. *b* is the slope of the xylem vulnerability curve. N is the number of simultaneous measurement points available for each species. C_a_ is the average atmospheric [CO_2_] during the dry-down experiment. Hydraulic traits ψ_50_ and *b* were preferably obtained from Martin-StPaul et al. (2017); traits obtained from Choat et al. (2012) are labelled with *; traits obtained using imputation methodology (Sanchez-Martinez et al., n.d.) are labelled with **.

| Species | Reference | Plant type / life form | $\psi_{50}$<br>[-MPa] | $b$<br>[-] | $N$ | $C_a$<br>[ppm] | Experiment<br>duration [days] |
| --- | --- | --- | --- | --- | --- | --- | --- |
| *Allocasuarina luehmannii* | (Posch & Bennett, 2009) | Evergreen sclerophyll angiosperm | 2.56 ** | 1.90 ** | 23 | 400 | 28 |
| *Glycine max* | (Liu et al., 2005) | Herbaceous | 1.90 ** | 1.43 ** | 9 | 380 | 21 |
| *Broussonetia papyrifera* | (Liu et al., 2010) | Deciduous broadleaf angiosperm | 0.49 * | 1.69 * | 6 | 360 | 120 |
| *Cinnamomum bodinieri* | (Liu et al., 2010) | Evergreen broadleaf angiosperm | 1.49 ** | 2.36 ** | 6 | 360 | 120 |
| *Platycarya longipes* | (Liu et al., 2010) | Evergreen broadleaf angiosperm | 1.25 * | 1.20 * | 6 | 360 | 120 |
| *Pteroceltis tatarinowii* | (Liu et al., 2010) | Evergreen broadleaf angiosperm | 0.93 * | 1.87 * | 6 | 360 | 120 |
| *Rosa cymosa* | (Liu et al., 2010) | Evergreen shrub | 2.26 ** | 1.88 ** | 9 | 360 | 120 |
| *Pyracantha fortuneana* | (Liu et al., 2010) | Evergreen shrub | 6.21 ** | 6.39 ** | 6 | 360 | 120 |
| *Ficus tikoua* | (Liu et al., 2011) | Evergreen shrub | 1.56 ** | 2.88 ** | 16 | 360 | 120 |
| *Cedrus atlantica* | (Grieu, Guehl & Aussenac, 1988) | Evergreen needleleaf gymnosperm | 4.98 | 5.05 | 9 | 300 | 35 |
| *Pseudotsuga menziesii* | (Grieu, Guehl & Aussenac, | Evergreen needleleaf gymnosperm | 4.82 | 5.68 | 7 | 300 | 35 |
|  | 1988) |  |  |  |  |  |  |
| *Pseudotsuga macrocarpa* | (Grieu, Guehl & Aussenac, 1988) | Evergreen needleleaf gymnosperm | 3.86 ** | 4.02 ** | 5 | 300 | 35 |
| *Cistus albidus* | (Galmés et al., 2007; Galmés, Medrano & Flexas, 2007) | Semi-deciduous sclerophyll shrub | 3.54 ** | 2.96 ** | 4 | 400 | 17 |
| *Hypericum balearicum* | (Galmés et al., 2007; Galmés, Medrano & Flexas, 2007) | Evergreen sclerophyll shrub | 2.36 ** | 1.93 ** | 4 | 400 | 17 |
| *Limonium gibertii* | (Galmés et al., 2007; Galmés, Medrano & Flexas, 2007) | Evergreen sclerophyll shrub | 1.79 ** | 1.31 ** | 4 | 400 | 17 |
| *Limonium magallufianum* | (Galmés et al., 2007; Galmés, Medrano & Flexas, 2007) | Evergreen sclerophyll shrub | 1.79 ** | 1.31 ** | 4 | 400 | 17 |
| *Malva subovata* | (Galmés et al., 2007; Galmés, Medrano & Flexas, 2007) | Semi-deciduous sclerophyll shrub | 1.58 ** | 3.61 ** | 4 | 400 | 17 |
| *Phlomis italica* | (Galmés et al., 2007; Galmés, Medrano & Flexas, 2007) | Semi-deciduous sclerophyll shrub | 4.51 ** | 2.35 ** | 4 | 400 | 17 |
| *Pistacia lentiscus* | (Galmés et al., 2007; Galmés, Medrano & Flexas, 2007) | Evergreen sclerophyll shrub | 6.41 ** | 8.28 ** | 4 | 400 | 17 |
| *Diploaxis ibicensis* | (Galmés et al., 2007; Galmés, Medrano & Flexas, 2007) | Herbaceous | 2.37 ** | 1.51 ** | 4 | 400 | 17 |
| *Beta maritima subsp. marcosii* | (Galmés et al., 2007; Galmés, Medrano & Flexas, 2007) | Herbaceous | 2.71 ** | 2.51 ** | 4 | 400 | 17 |
| *Beta maritima subsp. maritima* | (Galmés et al., 2007; Galmés, Medrano & Flexas, 2007) | Herbaceous | 2.71 ** | 2.51 ** | 4 | 400 | 17 |
| *Lysimachia minoricensis* | (Galmés et al., 2007; Galmés, Medrano & Flexas, 2007) | Herbaceous | 1.86 ** | 2.42 ** | 4 | 400 | 17 |
| *Eucalyptus pilularis* | (Kelly et al., 2016) | Evergreen sclerophyll angiosperm | 2.58 ** | 2.27 ** | 54 | 380 | 12 |
| *Eucalyptus populnea* | (Kelly et al., 2016) | Evergreen sclerophyll angiosperm | 5.64 * | 3.59 * | 65 | 380 | 12 |
| *Helianthus annuus* | (Tezara, Driscoll & Lawlor, 2008) | Herbaceous | 3.05 | 3.27 | 21 | 360 | 12 |
| *Olea europaea* var. *Chemlali* | (Ennajeh et al., 2008) | Evergreen sclerophyll angiosperm | 7.00 * | 2.07 * | 12 | 350 | 60 |
| *Olea europaea* var. *Meski* | (Ennajeh et al., 2008) | Evergreen sclerophyll angiosperm | 7.20 * | 2.45 * | 14 | 350 | 60 |
| *Picea abies* | (Salmon et al., 2020) | Evergreen needleleaf gymnosperm | 3.61 | 5.51 | 23 | 400 | 42 |
| *Pinus sylvestris* | (Salmon et al., 2020) | Evergreen needleleaf gymnosperm | 3.09 | 5.48 | 14 | 400 | 42 |
| *Betula pendula* | (Salmon et al., 2020) | Deciduous broadleaf angiosperm | 2.01 | 4.10 | 15 | 400 | 28 |
| *Populus tremula* | (Salmon et al., 2020) | Deciduous broadleaf angiosperm | 2.60 | 4.34 | 22 | 400 | 21 |
| *Quercus coccifera* | (Peguero-Pina et al., 2009) | Evergreen sclerophyll angiosperm | 6.00 ** | 9.01 ** | 6 | 350 | 12 |
| *Quercus suber* | (Peguero-Pina et al., 2009) | Evergreen sclerophyll angiosperm | 5.20 | 9.76 | 5 | 350 | 12 |
| *Quercus ilex* | (Peguero-Pina et al., 2009) | Evergreen sclerophyll angiosperm | 6.88 | 7.39 | 5 | 350 | 12 |
| *Quercus ilex* | (Epron & Dreyer, 1990) | Evergreen sclerophyll angiosperm | 6.88 | 7.39 | 14 | 350 | 20 |
| *Quercus petraea* | (Epron and Dreyer, 1990) | Deciduous broadleaf angiosperm | 3.30 | 4.30 | 19 | 350 | 20 |
| *Quercus pubescens* | (Epron and Dreyer, 1990) | Deciduous broadleaf angiosperm | 4.81 | 5.97 | 21 | 350 | 20 |
| *Sequoia sempervirens* | (Simonin, Santiago & Dawson, 2009) | Evergreen needleleaf gymnosperm | 5.66 ** | 4.28 ** | 17 | 400 | 21 |

### Species-specific model parametrisations and plant traits

All the stomatal optimization models studied here rely on a two-parameter hydraulic vulnerability curve (Eq. S4) to simulate water transport from roots to leaves. At the species-level, values of the water potential at 50% loss of xylem hydraulic conductance (ψ_50_) were obtained from hydraulic trait databases (preferably Martin-StPaul et al. (2017), otherwise Choat et al. (2012)). The slopes of the vulnerability curve (*b*) were estimated from (Eq. S4) and the water potential at 12% loss of hydraulic conductance (ψ_12_) or at 88% loss of hydraulic conductance (ψ_88_), accouting for the original equation used to describe the vulnerability curve. When species-level vulnerability parameters were not available (21 out of 38 species, Table 1), we performed an imputation of the missing values using environmental, phylogenetic, and trait correlations and covariations. These correlations and covariations derived from the entire Xylem Functional Traits Database (Choat et al., 2012) alongside environmental data (mean annual temperature and average annual precipitation) extracted from the CHELSA database (Karger et al., 2017), following the methodology developed by Sanchez-Martinez et al. (n.d.). The relationships obtained in this way were used to estimate species-specific values of ψ_50_ and *b*.

Another key parameter of the stomatal optimization models is K_max_ (mol m^-2^ s^-1^ MPa^-1^; Eq. S3), which sets the maximum conductance to water flow between the roots and the leaves. The estimates of K_max_ available from the literature are seldom suitable for modelling efforts (because models require parametrisations of K_max_ at the scale of the whole plant, measured per unit leaf area; Mencuccini, Manzoni & Christoffersen, 2019). Therefore, here, we calibrated K_max_ for each species within each stomatal optimization model and dry-down experiment (see the ‘Parameter calibrations and simulation experiments’ section below).

Finally, to analyse the potential coordination between α and other plant traits (objective 3), we collected species-specific traits related to water use strategies and plant form and function. These were the species’ maximum height (H_max_; m), specific leaf area (SLA; cm^2^ g^-1^), Huber value (Hv; i.e., sapwood area per leaf area in cm_sw_^2^ m_leaf_^-2^), and maximum leaf-area specific hydraulic conductivity (*K_L_*; mol m_leaf_^-1^ s^-1^ MPa^-1^), all of which were retrieved from the Xylem Functional Traits Database (Choat et al., 2012). Missing species-specific trait values were obtained by imputation (as described above for ψ_50_ and *b)*, following Sanchez-Martinez et al. (n.d.) (See Table S1 for the RMSE of each imputed variable).

### Parameter calibrations and simulation experiments

We conducted a different set of parameter calibrations for each of the three objectives of this study (Fig. 1). Our first set of calibrations was designed to explore the co-variation of α with hydraulic (or hydraulic cost) parameters in different models, as well as its relation to other functional traits and environmental conditions (Objective 3). To obtain the best estimates of α, we fitted all model parameters that influence the optimality criteria. By contrast, our second set of parameter calibrations was much more conservative, as it aimed to infer the models’ performance gain from photosynthetic acclimation in the absence of a species-specific knowledge of α (Objective 1). To that end, we fitted one hydraulic (or hydraulic cost) parameter per model while keeping all other parameters fixed. Our third set of parameter calibrations was used to gain insight into the relative contributions of hydraulic (stomatal) vs. non-stomatal limitations on photosynthesis (Objective 2). In this case, we fitted two model parameters: one hydraulic (or hydraulic cost) parameter and α.

**Figure 1.**
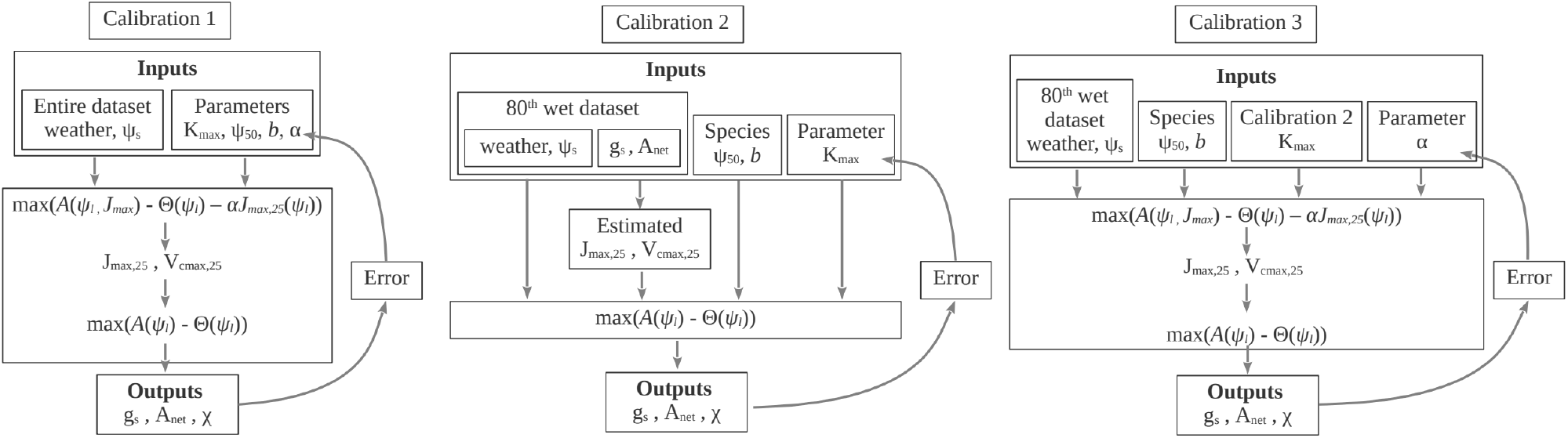
Schematics of the conducted parameters calibrations for the models PMAX, PMAX2, and SOX. Models with an additional hydraulic cost parameter (i.e. SOX2, PHYDRO, and CGAIN) would also have this parameter calibrated during the first calibration. In calibration 2, instead of calibrating the K_max_, of these models, their specific hydraulic parameter was calibrated, and the K_max_ parameter calibrated using the PMAX model was used. Parameter optimization was achieved by minimizing the error between predicted and measured instantaneous g_s_, A_net_ and C_i_:C_a_ ratios using Eq. 10. The parameters box shows the parameters calibrated in each set of calibrations.

In our first set of calibrations (“complete calibration”), we accounted for photosynthetic acclimation as described in the ‘Acclimation of the photosynthetic capacity’ section (Eq. 9). Using the Nelder-Mead algorithm, we looked to fit the best {K_max_, ψ_50_, *b*, α} parameter set for each individual model, species, and dry-down experiment that minimized the normalised root mean square error (NRMSE) between predicted and measured instantaneous rates of g_s_, A_net_ and C_i_:C_a_ ratios. Compared to the PMAX, PMAX2, PMAX3, and SOX models which only required calibrating the aforementioned parameters, the SOX2, PHYDRO, and CGAIN models required an additional hydraulic cost parameter to be calibrated (i.e., δ, γ, and ϖ, respectively).

The first set of calibrations was performed on the entire dataset of each species, and it was also repeated without photosynthetic acclimation (Eq. 1). Calibrating four or five parameters on small data subsets (e.g., N=4; Table 1) likely leads to unrealistic predictions of these parameters. Therefore, the parameter estimates from small data samples were given less weight in subsequent statistical analyses.

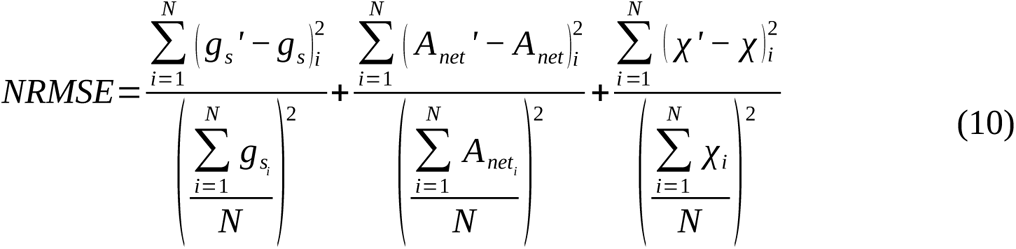

where *χ* is the ratio of C_i_:C_a_, the prime symbol ‘ denotes predicted values, the double-subscript *i* represents the position of the paired predicted and actual values, and N is the total number of actual observations for a given species in an experiment.

We extracted the average α of all species from the first set of calibrations, considering the respective biases of each stomatal model and data subset by fitting a crossed random intercept of stomatal models, species and dry-down experiments in a mixed-effects linear model (average α = 0.0945). Setting α to the average of all species and dry-down experiments, and with species-specific values of ψ_50_ and *b* retrieved from the literature (Table 1), we conducted our second set of calibrations. In the PHYDRO model, instead of using the parameters of the xylem vulnerability curve, we used an approximate estimate of the outside-xylem (ox) vulnerability curve by setting ψ_ox50_ = ψ_50_/3 and *b*=1, based on the findings of Joshi et al. (2022). We calibrated K_max_ (or the model-specific parameters of SOX2, PHYDRO, and CGAIN using the K_max_ parameter obtained from PMAX) on the 80^th^ wet percentile data of each species and dry-down experiment, using a successive parabolic interpolation algorithm and the parameter fitting criterion given by Eq. 10. These parameter calibrations can be deemed “average α acclimated”.

Our last set of parameter calibrations (“calibrated α acclimated”) prescribed the hydraulic parameter values obtained in the second set of calibrations, and α was calibrated on the 80^th^ wet percentile data of the species and dry-down experiment using the same parameter fitting criterion given by Eq. 10.

Both acclimated (Eq. 9) and non-acclimated (Eq. 1) subsequent model simulations were performed mostly outside their calibration samples, over the entire dry-down experiments data for a given species-experiment. We opted to use the hydraulic (or hydraulic cost) parameters obtained for the non-acclimated models when running their acclimated counterparts, in order to make the simulation outcomes as comparable as possible. In all calibrations and model simulations without acclimation, J_max,25_ and V_cmax,25_ were obtained using the average values of the actual g_s_ and A_net_ measurements for the 80^th^ wet percentile of each species-study (Fig. S1-S2), and assuming photosynthetic coordination via Eqs. S6, S11, S12, and S13.

Finally, to ensure that using a different parameterisation of hydraulic vulnerability did not give the PHYDRO model an unfair advantage or disadvantage compared to other models, we repeated the second and third set of calibrations (and associated simulations) using outside-xylem vulnerability curve parameters (i.e., ψ_ox50_ = ψ_50_/3 and *b*=1) for other models and xylem vulnerability curve parameters (i.e., ψ_50_ and *b*) for PHYDRO. This incidentally provided us with a test of the sensitivity of the models to the choice of vulnerability curve parameters.

### Statistical analyses

The performance of the seven stomatal models in predicting A_net_ with and without acclimation (Objective 1) was assessed using four statistical metrics computed at the level of each species in each dry-down experiment: (1) Pearson’s correlation coefficient, where coefficients close to 1 indicate a perfect positive correlation; values ≤ 0 indicate a null or negative correlation, (2) The slope (*m*) between the estimated and actual A_net_ measures the proportional bias of a model’s predictions. Values of *m* close to 1 indicate good proportionality between the estimated and actual A_net_, whereas *m* much smaller/higher than 1 imply that the predicted A_net_ are underestimated/overestimated as the soils dry out, (3) RMSE, which measures the accumulated error of A_net_ predictions, with a larger penalty on larger errors. The lower the RMSE, the lower the accumulated error, and (4) bias (Eq. 11) of the A_net_ estimates evaluates the average simulation error and its direction. A negative bias implies that the estimated values are, on average, systematically higher than the actual values.

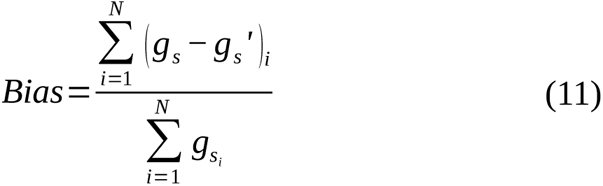

For each statistical metric, we used linear mixed-effects models (LMMs) to further compare the performance of models parametrised with the second set of calibrations (i.e., species-specific values of ψ_50_ and *b,* average α of all species and dry-down experiments). The models, with or without acclimation and their interactions were used as fixed explanatory variables. The dry-down experiments and species were used as crossed random intercepts to account for potential systematic errors in the experimental design or potential species-specific biases that could be introduced across stomatal models. All performance analyses were repeated for g_s_.

We used model with or without acclimation parametrised according to our third set of calibrations, to evaluate how each stomatal optimisation model divides its predictive capacity into the components of hydraulic impairment vs. limitations due to photosynthetic acclimation (Objective 2). For each stomatal model, we analysed how the explained variability (R^2^) of A_net_ was partitioned between the hydraulic penalty (stomatal partition), the cost of maintaining photosynthetic capacity (αJ_max,25_; non-stomatal partition), and species. We accounted for the explained variability by species because species-level ψ_50_ and *b* parameters may be inadequate for a given population within the species. The stomatal variance partition was computed as the marginal R^2^ of the actual A_net_ explained by the estimated A_net_ without acclimation and adjusting random intercepts for each species in an LMM. We then applied this LMM to the estimated A_net_ values with acclimation, and we obtained the non-stomatal variance partition as the difference between its marginal R^2^ and the previously calculated marginal R^2^ for the stomatal variance partition. The species variance partition was obtained as the difference between the conditional and the marginal R^2^ of the LMM for the acclimated A_net_ simulations. To understand whether the importance of these variance components changes as the soil dries, we repeated the analysis on the driest 50^th^ percentile of data for each species within each dry-down experiment.

Finally, we analysed the relationships between the calibrated α from the first set of calibrations and other species– and experiment-specific traits and growing conditions. The traits considered as possible explanatory variables for α were the species-level H_max_, ψ_50_, SLA, Hv, and K_L_ collected or imputed from the literature (Table 1). We selected these traits as they stand for different axes of variation representing different water use strategies and life forms. We also included an estimate of the actual J_max,25_ under well-watered conditions (80^th^ wet percentile; J_maxWW,25_), which was inferred from the observed A_net_ and g_s_ using the one-point method (Eqs. S6, S11, S12, and S13). The environmental conditions of interest were average I_abs_ and C_a_ during the experimental period.

During the experiments, I_abs_ was strongly positively related to VPD and air temperature (Fig. S3). All explanatory variables that presented a non-normal distribution (H_max_, SLA, Hv, K_L_, and J_maxWW,25_) were transformed using the natural logarithm to minimize the leverage effect of large values. We performed a multivariate LMM in which the variability of α was explained by the chosen species-level traits, ψ_50_, J_maxWW_, and the growing conditions, with the stomatal models and dry-down experiments used as crossed random intercepts. Because species-level traits were taken as fixed predictors of the LMM, it was important we used the stomatal models as random intercepts to account for the variability in α caused by each stomatal model for each species. Next, we identified the most significant predictors using an Akaike Information Criterion (AIC) stepwise algorithm.

Finally, we calculated the variance inflation factor (VIF) of each predictor to explore possible multicollinearity across explanatory variables. We also repeated the above model and the identification of the most significant predictors using the product of α and J_maxWW,25_, which corresponds to a photosynthetic capacity cost under well-watered conditions, as an explanatory variable.

In all LMMs, we used the natural logarithm of each species’ sample size as a weighting factor to minimize the effect of a small number of data points (see N, Table 1).

### Code

All analyses and calibrations were performed using R 4.2.0 (R Core Team, 2022) and the code is freely available from GitHub at https://github.com/vflo/Acclimated_gs_optimization_models. The following specific R packages were used:

- TrEvol (Sanchez-Martinez et al., n.d.) for the imputation of missing species-level trait values (see the ‘Species-specific model parametrisations and plant traits’ section above);
- dEoptim (Mullen et al., 2011) to solve the instantaneous optimality criteria presented in Eqs. 1 and 2, and 9 and 10;
- lme4 (Bates et al., 2015) for all LMM computations (see ‘Statistical analyses’);
- MuMIn (Bartoń, 2020) for the R^2^ variance partitions (‘Statistical analyses’).

## Results

All three calibration sets were successful for all the species and stomatal models. Most of the model parameterisations obtained from the first set of calibrations were insensitive to acclimation (Figs. S4a, S4b and S4c), with the exception of the magnitude of –ψ_50_ and K_max_ from SOX and SOX2 which were reduced by acclimation. In the second set of calibrations, where K_max_ was the only calibrated parameter for PMAX, PMAX2, PMAX3 and SOX, PMAX and SOX’s K_max_ were much lower than those from the first set of calibrations in the absence of acclimation (Fig. S4a), and moderately lower for PMAX2 and PMAX3 (p<0.05). For PMAX, K_max_ also substantially decreased between the first and second sets of parameter calibrations accounting for acclimation (Fig. S4a).

### Model performance with vs. without photosynthetic acclimation

All seven stomatal models generated highly and positively correlated in-sample A_net_ estimates across most species (Fig. 2a, Fig. S5). The mean Pearson’s correlation coefficient for the stomatal models without acclimation ranged from 0.761 in PMAX2 to 0.812 in PHYDRO, whilst for the acclimated stomatal models using the average α, it ranged between 0.804 and 0.823 in PMAX2 and PMAX, respectively, and using the calibrated α, it ranged from 0.794 in SOX to 0.817 in PMAX (Fig. 2a). The inclusion of acclimation produced a moderately positive effect on the estimates of PMAX2 and PMAX3 in terms of correlation, whereas the effect was negligible for CGAIN, PHYDRO, PMAX, SOX, and SOX2 (Fig. 2a).

**Figure 2.**
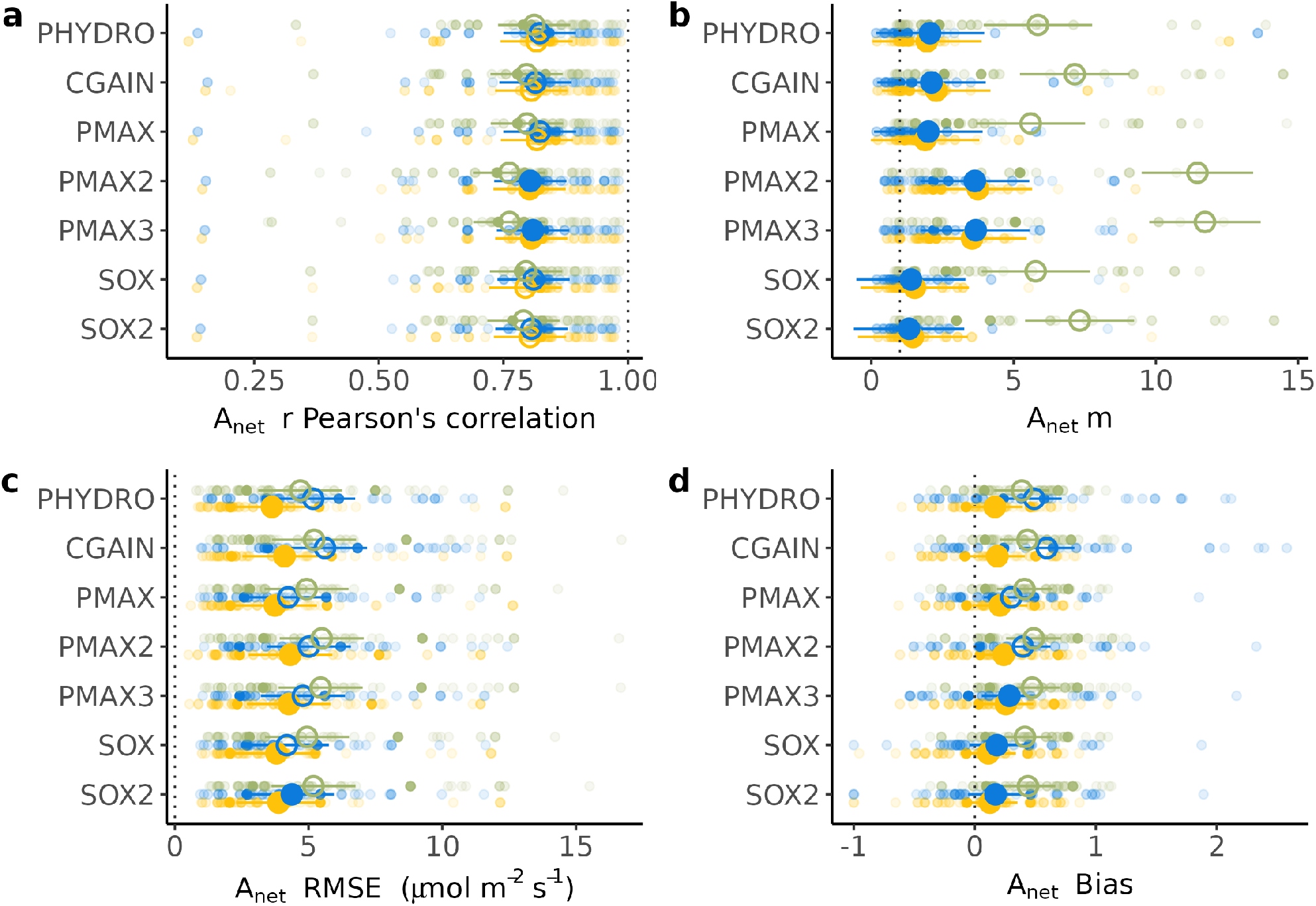
Performance comparison between acclimated and non-acclimated stomatal optimisation models, relative to species-specific observations of A_net_ during different dry-down experiments. All the performance metrics were calculated for estimates of A_net_ obtained within their full data sample, having fitted just one hydraulic parameter per stomatal model on the 80^th^ wet percentile of the species data samples (not acclimated), then acclimating by setting α to 0.0945 for all the species (average α), and finally acclimating with a calibrated α. For all the models except PHYDRO we used the species-level vulnerability curve parameters ψ_50_ and *b* shown in Table 1. For PHYDRO we used species-level outside-xylem vulnerability curve parameters as ψ_ox50_ = ψ_50_/3 and *b* = 1. Small circles are the values for each species within each dry-down experiment, higher transparency indicates smaller data sample. Large circles are the metric averages estimated using an LMM (see Methods). Large closed circles indicate a significant difference in the metric averages of the model realisations with acclimation with respect to not acclimated ones; large open circles indicate no significant difference. Statistical significance was calculated using paired t-test comparisons. Vertical dotted lines indicate the best achievable performance for each metric. a) Pearson’s correlation b) slope (*m*) of the linear relationship between the actual and estimated A_net_ c) RMSE d) bias of the estimation of A_net_ calculated as in Eq. 11.

By contrast, the slope between the predicted and actual A_net_ values (*m*) was strongly influenced by acclimation. *m* was significantly larger when acclimation was not accounted for (Fig. 2b) because the models failed to simulate the downregulation of A_net_ as the soil dried out in the absence of acclimation. Upon acclimation, the average values of *m* for CGAIN, PHYDRO, PMAX, SOX, and SOX2 were not likely to differ from one (H_1_ ≠ 1; P>0.05), indicating that these five models’ estimates of A_net_ were in good proportionality to the observations as the soil dried. The average *m* of PMAX2 and PMAX3 was greater than one though, most probably because they failed in a few species-experiments with a large N (Fig. 2b).

The different models did not significantly differ from one another in their average non-acclimated RMSE. However, acclimated models with species-experiment specific α (i..e, calibrated α acclimated from the third set of calibrations), all displayed a significantly lower RMSE than those without acclimation (Fig. 2c). In addition, the average α acclimated (second set of parameter calibrations) SOX2 had significantly lower RMSE than without acclimation, and the average α acclimated CGAIN had significantly higher RMSE than PMAX, SOX and SOX2 (Table S2). The calibrated α acclimated PHYDRO presented the lowest average RMSE for A_net_ at 3.63 μmol_C_ m^-2^ s^-1^ (Fig. 2c, Table S2), whereas the average α acclimated CGAIN presented the highest average RMSE at 5.61 μmol_C_ m^-2^ s^-1^.

All the calibrated α acclimated models presented biases significantly lower than the non-acclimated models and not significantly different to zero, with the exception of the calibrated α acclimated PMAX2 and PMAX3 for which biases were slightly lagger than zero (p<0.05). PHYDRO, CGAIN, PMAX and PMAX2 did not show significant differences in biases between their average α acclimated and their non-acclimated model versions (Fig. 2d). All the non-acclimated stomatal models presented a significant average overestimation (bias higher than zero; p<0.05), with PMAX2 presenting the larger bias (0.486) and PHYDRO the lowest one (0.388; Fig. 2d).

Unlike for A_net_, the models’ performance for g_s_ in terms of Pearson’s correlation, *m*, RMSE, and bias was generally not affected by the inclusion of acclimation (Fig. S7). Yet, acclimation led to a significant improvement of proportionality for CGAIN, PMAX2 and PMAX3 (Fig. S7b). However, *m* was still significantly higher than 1 for PMAX2 and PMAX3, pointing to these models’ failure to produce estimates of g_s_ in good proportionality with the observations (Fig. S7b).

Importantly, all models except the PMAX variety were highly sensitive to different vulnerability curve parameters (Fig. S6), poor predictions result when parameters other than those prescribed in the respective model designs were used. Thus, using the outside-xylem vulnerability curve parameters, both SOX and SOX2 exhibited enhanced sensitivity to changes in ψ_s_, leading to rapid and strict stomatal closure as soil dried (Fig. S6b and S6d). CGAIN became unstable, and for several species, the model failed to converge. By contrast, both PMAX2 and PMAX3 showed improved performance (Fig. S6) and similar estimates to PMAX. Likewise, using the xylem-level vulnerability curve parameters, PHYDRO was unable to accurately depict the decline of A_net_ as the soil dried out (Fig. S6b).

### Hydraulic impairment vs. limitations due to photosynthetic acclimation

The proportion of variance in A_net_ explained by hydraulic (stomatal limitation) and metabolic impairment (non-stomatal limitation to photosynthesis) differed among models (Fig. 3). Overall, stomatal limitations were markedly more important than non-stomatal limitations for all models across all conditions (Fig. 3a), except for PMAX2 and PMAX3 in which non-stomatal limitations equally mattered. When analysing the driest half of the data, the variance explained by non-stomatal limitations slightly decreased for all models (Fig. 3b). The acclimated PMAX model explained the greatest amount of variability in A_net_, with a marginal R^2^ of 0.59 for all data points (stomatal + non-stomatal contributions), whereas the acclimated PMAX, PMAX2 and PMAX3 were the best models for the driest half of the data, achieving a marginal R^2^ of 0.55. The proportion of variance explained by the different species was greater for the driest half of the data (Fig. 3a), accounting for nearly half of the variance explained by the stomatal and non-stomatal components taken together, which hints at site-specific adaptations or acclimations and at increased errors introduced by species-level parameterisations when water supply is limiting.

**Figure 3.**
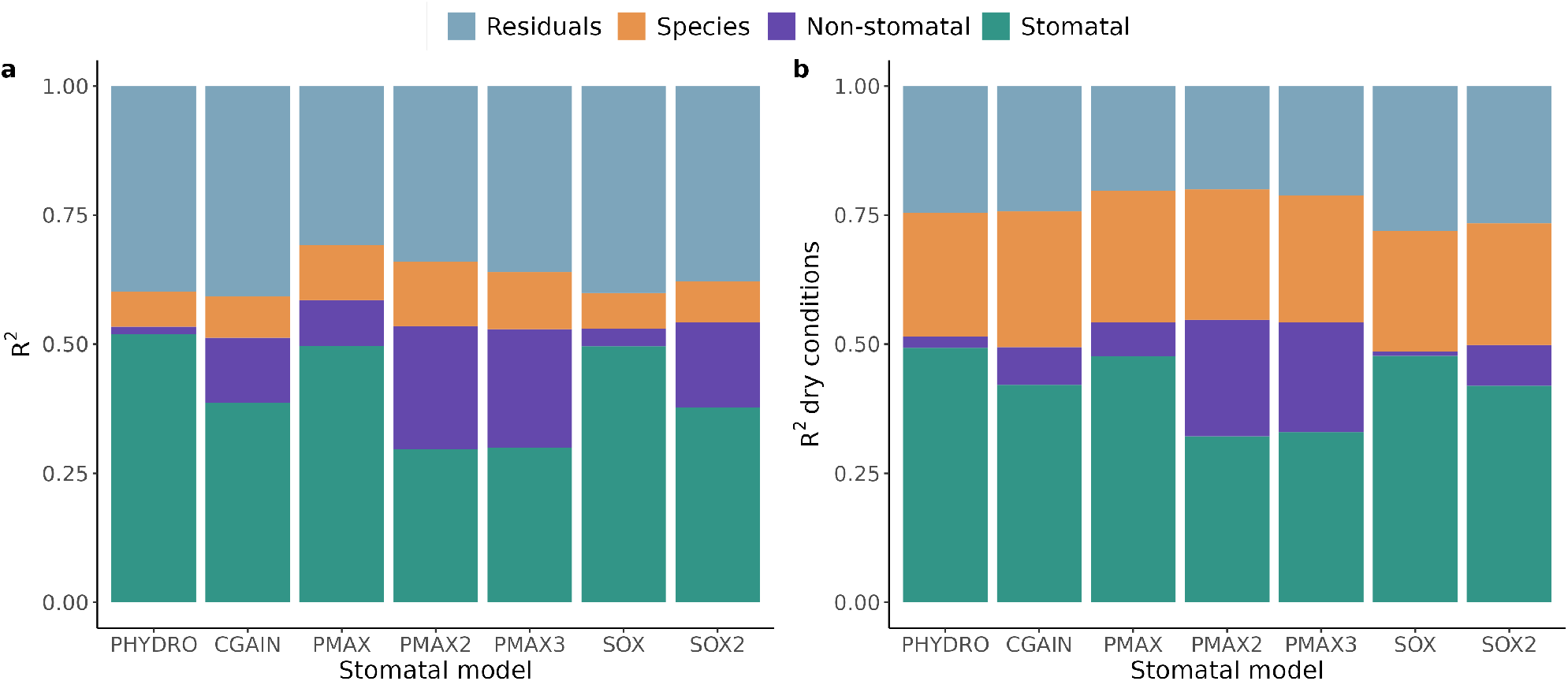
Portion of A_net_ variance explained by stomatal, non-stomatal, and species-level contributions for each stomatal model. a) partition using all the data b) partition using the driest 50^th^ percentile data for each species. Grey portions represent any residual variance not explained by stomatal, non-stomatal, or species-level contributions.

### Relationship between α, functional traits, and environmental growing conditions

Calibrating {K_max_, ψ_50_, *b*, α} (and additionally, γ, δ or ϖ, for PHYDRO, SOX2, and CGAIN, respectively) for each model, species, and dry-down experiment (first set of parameter calibrations) enabled a majority of the models to match or exceed a R^2^ of 0.6 for both g_s_ and A_net_ (Table S3); PMAX2 and PMAX3 were exceptions in that they only exceeded 0.5 for g_s_. These high R^2^ values gave us confidence in the average α obtained both across models and for each model. The PHYDRO and CGAIN models, which require an additional hydraulic cost parameter, exhibited a significantly higher average α (α > 0.1024) than other models, all of which exhibited significantly different average α from one another (α between 0.087 – 0.0942; Fig. 4). Despite these differences, none of the model’s average α was different from the average α across models (α = 0.0945) nor different from 0.1, the value proposed by Joshi et al. 2022 for global applications.

**Figure 4.**
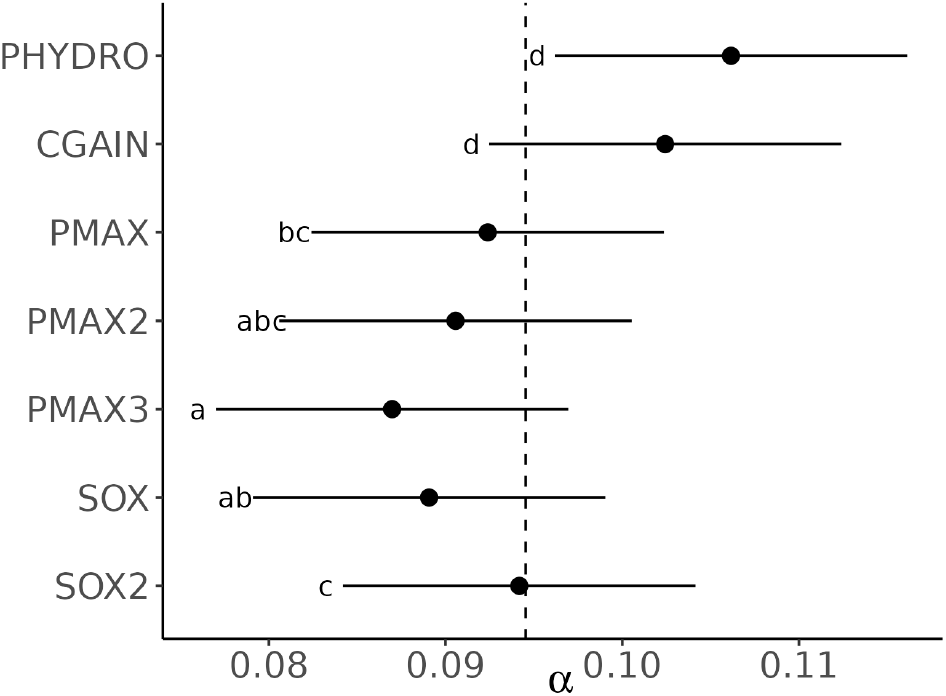
Average model-specific estimate of α across species, accounting for the random effects of the species and dry-down experiments. Vertical dashed line show the average α across models (α = 0.0945). Letters represent groups with significant differences calculated using Sidak’s multiple comparison test.

A multivariate LMM combined with a stepwise AIC variable selection (Table 2) revealed three plant traits and growing conditions to explain a moderate part of the variability in α across species (marginal R^2^ = 0.47), out of the eight potential explanatory variables originally considered. The species maximum height and J_maxWW,25_ (i.e., the J_max,25_ inferred from observations under well-watered conditions) were negatively related to α, whilst the average I_abs_ had a positive relationship with α (Table 2, Fig. 5). Multicollinearity was low across the explanatory variables (VIF; Table 2).

**Figure 5.**
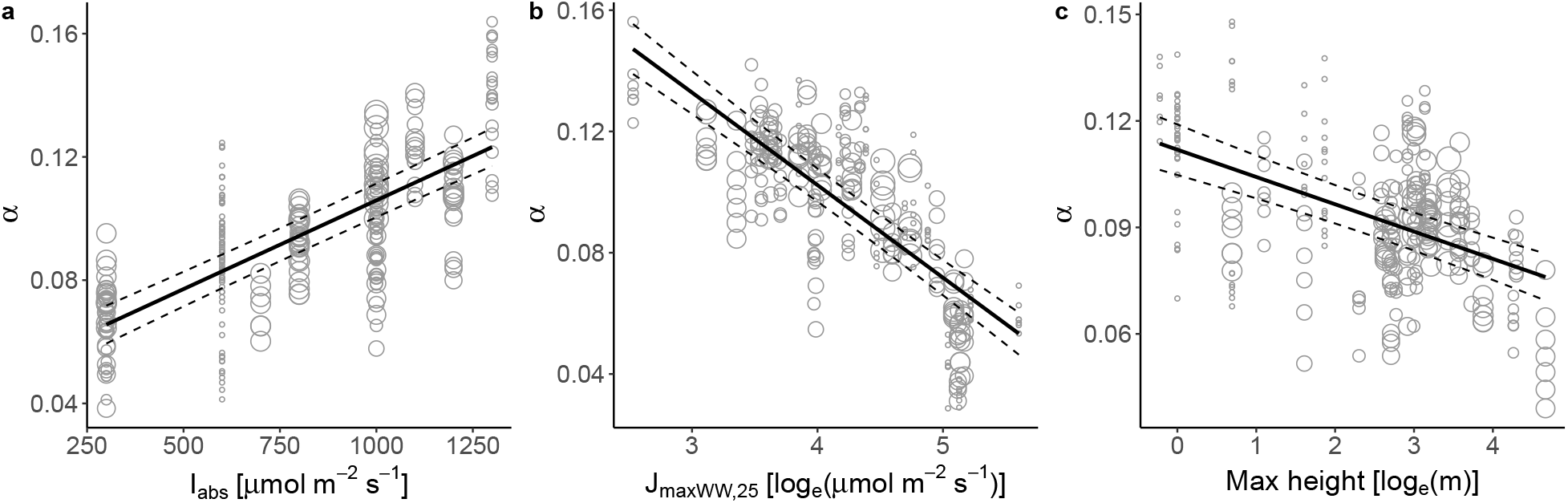
Partial effects from a multivariate Linear-Mixed Model (LMM) explaining the variability of α. The LMM was fitted using the dry-down experiments and stomatal models as crossed random intercepts and the LMM coefficients are shown in Table 2. Grey scatter open points show all the model-specific contributions to α, across all species. Black plain lines are partial regressions and dashed lines represent the partial regressions plus or minus their standard error. Size of the points represent the weight applied to each data point, which is equal to the natural logarithm of the sample size (Table 1).

**Table 2.** Multivariate linear-mixed effects model (LMM) best explaining the variability of α for the fully calibrated stomatal optimization models (first calibration set from the methods). Iabs is the average absorbed photosynthetic active flux radiation during the experiment in μmol m^-2^ s^-1^. J_maxWW,25_ is the maximum electron transport capacity at standard temperature (25°C) inferred from observations under well-watered conditions in log_e_(μmol_C_ m^-2^ s^-1^). H_max_ is the species maximum height in m. The Estimate, SE, and VIF columns stand for the estimated coefficients, standard error, and variance inflation factor.

|  | Estimate | SE | VIF | T value | p-value |
| --- | --- | --- | --- | --- | --- |
| Intercept | $1.96 \times 10^{-1}$ | $9.50 \times 10^{-2}$ | | 20.67 | <0.001 |
| Iabs | $5.76 \times 10^{-2}$ | $3.10 \times 10^{-3}$ | 1.04 | 18.56 | <0.001 |
| $J_{\max WW,25}$ | $-3.05 \times 10^{-2}$ | $1.78 \times 10^{-3}$ | 1.57 | -17.17 | <0.001 |
| Hmax | $-7.69 \times 10^{-3}$ | $9.47 \times 10^{-4}$ | 1.57 | -8.12 | <0.001 |

The results were similar albeit explaining a larger part of the variability (R^2^ = 0.59) when the relationships between αJ_maxWW,25_ and the eight potential explanatory variables were explored (Fig. S8). In that latter case, six explanatory variables were selected: the SLA and maximum height of the species were associated with lower photosynthetic capacity cost under well-watered conditions, whilst I_abs_, species K_L_, H_v_, and ψ_50_ were positively related to αJ_maxWW,25_ (Fig. S8).

## Discussion

This study demonstrates that accounting for a maintenance cost for photosynthetic capacity, as proposed by Joshi et al. (2022), improves predictions of A_net_ (Fig. 2) across a range of hydraulics-enabled stomatal optimality frameworks, particularly under drought conditions (Fig. 2b). Our approach eliminates the need for species-level knowledge of photosynthetic capacity (e.g., J_max25_ and V_cmax25_), for empirical functions describing the sensitivity of photosynthetic capacity to drought (Drake, 2017; Egea et al., 2011; Keenan et al., 2009; Knauer et al., 2019; Yang et al., 2019; Zhou et al., 2014, 2013), or for semi-empirical functions relating photosynthetic capacity to leaf nitrogen investments (Sabot et al. 2022a). Despite these benefits, our approach requires fitting an additional photosynthetic cost parameter α, whose variable responses to species-specific plant traits and current environmental conditions (let alone future conditions) have not been fully elucidated in this study. Nonetheless, our results suggest that the cost parameter α could be predicted from plant functional traits and parameters, in combination with knowledge of the relevant growth conditions.

Despite variability in model-specific α values (Fig. 3), the predictions of the seven stomatal models taken together allowed us to meaningfully explore the relationships between α and other plant traits and environmental conditions, irrespective of the models’ hydraulic impairment functions. Our results revealed a strong relationship between α and J_maxWW,25_, implying that plants with lower photosynthetic costs under well-watered conditions have higher levels of photosynthetic capacity.

Whether photosynthetic costs change through time or increase during dry-downs is not resolved by our analysis, but it is plausible that the magnitude of α might depend on hydraulic impairment. Interestingly, we found a positive relationship between α and I_abs_ which suggests that species experiencing higher light intensities have higher leaf maintenance costs per unit J_max25_. This seemingly contradicts our expectation, as species growing under higher radiation levels are typically characterized by higher J_max25_ (Peng et al., 2021). However, there is also evidence of more pronounced photo-inhibition for leaves under high light intensities exposed to moderate to severe water deficits, compared to leaves under low radiation (Kaiser, 1987; Epron & Dreyer, 1990). This suggests an enhanced photosynthetic sensitivity to drought (i.e., a higher α) for high J_max25_ leaves, perhaps linked to the production costs of photoprotective compounds whose concentrations acclimate to UV radiation levels (Barnes et al., 2023). Further considering the strong positive correlation between I_abs_ and VPD (Fig S3), it is likely the increased cost associated with elevated radiation is influenced by the confounding effect of higher atmospheric aridity.

Besides its relations to environmental drivers, α was negatively related with species maximum height but positively related with ψ_50_ and K_L_ (Fig. S8). This suggests a coordination of the photosynthetic cost with the hydraulic safety-efficiency trade-off, as well as with the life form strategy. Such a pattern would agree with the observational evidence that species with higher K_L_ have smaller hydroscapes and close their stomata at less negative water potentials (Fu et al., 2019; Fu & Meinzer, 2019; Santiago et al., 2004; Li et al., 2019), hence, in the context of this study, requiring a higher α to prompt a rapid decrease in photosynthetic capacity as the soil dries out. In the same vein, taller species typically show more ‘efficient’ water use strategies (Flo et al., 2021) and coordination with ψ_50_ and K_L_ across species and life forms (Liu et al., 2019). This coordination implies that taller plants would typically have stronger stomatal regulation as the soil dries out, and therefore a higher photosynthetic cost may be anticipated. However, note that all woody plants included in this study were seedlings, and many of them were far from their maximum potential heights. Nevertheless, the maximum height of the species still successfully represents the axis of variation of the different life form strategies.

With the caveat that performance evaluation of the models presented here only accounts for model predictions of A_net_ and g_s_, PMAX exhibited the best performance when photosynthetic acclimation was included (Fig. 2 and 3). Previous studies have shown this stomatal optimisation model to perform as well or better than empirical stomatal models in the absence of photosynthetic acclimation (Sabot et al., 2022b; Venturas et al., 2018; Wang et al., 2020), and it has also been shown to make robust predictions when accounting for it (Sabot et al. 2022a). The SOX, SOX2, PHYDRO, and CGAIN also exhibited reasonably good performance (Fig. 2 and 3), especially in terms of improvement to proportionality when acclimation was considered (Fig. 2b), but note that these four models were more sensitive to the choice of vulnerability curve parameters than PMAX (cf. Fig. 2 and S6). In stark contrast, even though the inclusion of photosynthetic acclimation improved the performance of PMAX2 and PMAX3 in terms of correlation, RMSE, and bias, it did not improve the proportionality of their A_net_ predictions (Fig. 2b) except when outside-xylem vulnerability curve parameters were used (Fig. S6). Overall, our simulations’ performance without acclimation did not support the assertion made by Wang et al. (2020) that PMAX2 outperforms other stomatal optimisation models, in line with the findings of Sabot et al. (2022b).

Stomatal optimisation accounting for hydraulic impairment functions alone demonstrated enough flexibility to predict A_net_ reasonably well when using species-level prescribed parameters of vulnerability curves and calibrating other hydraulic parameters (Table S2, “not acclimated”), particularly for those models that incorporate an additional hydraulic parameter besides just K_max_ (i.e., CGAIN, PHYDRO, and SOX2). However, all stomatal optimization models examined here except one (PMAX) were sensitive to the choice of hydraulic vulnerability curve parameterisation. Therefore, they might not perform so well when directly parameterised based on both hydraulic vulnerability curves and estimates of K_max_ for a given plant organ (e.g., leaf, branch, stem, root), especially if those estimates are taken in isolation from parameters related to photosynthetic acclimation, which we have shown to positively affect model outcomes. The difficulty lies in determining how the vulnerability of a plant organ should be scaled to better represent the vulnerability of the entire hydraulic system for each model – this is complex because organ-level traits do not accurately characterize whole-plant hydraulic responses (Wang and Frankenberg 2022) – and integrated with our understanding of other plant functions – this is complex because our understanding of trait coordination at the plant level is incomplete (McCulloh et al., 2019). We recommend strong caution and stress the need for in-depth testing before the application of stomatal optimisation models at large spatial scales, let alone before their integration into global vegetation models and land surface models.

## Acknowledgements

The authors would like to express their gratitude to all the contributors to the research from which the experimental data were drawn, and especially Jaume Flexas and Jeroni Galmes for sharing their original database. Additionally, the authors extend their thanks to Shuangxi Zhou, who compiled the first version of the database used in this study, to Rafael Poyatos for providing access to computing facilities, and to Rodolfo Nobrega for the key discussions at an early stage of the study. Victor Flo acknowledge the Spanish Ministry of Universities who supported him through a Margarita Salas grant (698511). Manon Sabot acknowledges support from the Australian Research Council (ARC) Centre of Excellence for Climate Extremes (CE170100023), as well as the ARC Discovery Grant (DP190101823). Jaideep Joshi gratefully acknowledges funding from the European Union’s Horizon 2020 research and innovation programme under the Marie Skłodowska-Curie Actions fellowship (grant agreement No. 841283 – Plant-FATE), the Strategic Initiatives Program of the International Institute for Applied Systems Analysis (IIASA; Project name – RESIST), and the National Member Organizations that support IIASA. DS and ICP acknowledge support from the European Research Council (787203 REALM) under the European Union’s Horizon 2020380 research programme.

## Contributions

VF conceptualized the study with the help of all authors. VF and JJ prepared the database. VF and JJ prepared the code. MS and DS supervised the code. VF conducted the calibrations, simulations, and statistical analyses. VF wrote the first draft. All the authors provided scientific input, and reviewed and edited the final manuscript.

## Notes

https://github.com/vflo/Acclimated_gs_optimization_models

### Competing Interest Statement

The authors have declared no competing interest.

