## Supplementary Material for "Incorporating photosynthetic acclimation improves stomatal optimisation models"

##### Notes 1

Certain elements of the theory outlined in this section has been extensively covered in other works (Joshi et al., 2022; Wang et al., 2017) but are presented here for convenience.

Water transport model

The hydraulic modelling framework we use is based on a simple water transport model at steady-state (Sperry et al., 2016). Under stationary conditions, the volumetric water transpired by the plant (Eq. S1) matches the volumetric water supplied by the soil (Eq. S2). This balance maintains the water continuum across the soil-plant-atmosphere and thus prevents embolism in the conductive tissues. The transpiration stream is hence adjusted to particular conditions of atmospheric evaporative demand and soil water potential by regulating the stomatal conductance ($g_s$). Stomatal regulation is simultaneously linked to the leaf water potential ($\psi_l$) and thus determines the water potential differential between the soil and the leaf ($\Delta\psi$), which, together with the characteristics of the roots and conductive tissues, governs the water supply.

Transpiration per unit leaf area can be approximated using Flick's law of diffusion,

$$E = 1.6 g_s (e_l - e_a) \approx 1.6 g_s D \qquad \text{(S1)}$$

here, $g_s$ is the stomatal conductance to $CO_2$, the 1.6 factor converts from conductance to $CO_2$ to conductance to water vapour, $e_l$ and $e_a$ are the relative partial pressure of water vapour inside the leaf near the stomata cavities and in the atmosphere, respectively, and D is the water vapour deficit of the atmosphere. For the right-hand side approximation of the equation, we assume that $e_l$ is always saturated, so that is about the atmospheric partial pressure saturation (i.e. $e_l \approx e_s$) and therefore $e_l - e_a \approx e_s - e_a = D$.

The supply function is based on Darcy's law, were the volumetric flow of water (Q) is given by the conductance of the entire water pathway and by the pressure drop from the soil ($\psi_s$) to the leaves ($\psi_l$).

$$Q=\int_{\psi_s}^{\psi_l} k_\psi \, d\psi \qquad \text{(S2)}$$

where $k_\psi$ is the hydraulic conductance at a given water potential and is defined as,

$$k_\psi = K_{max} P(\psi) \qquad \text{(S3)}$$

where $K_{max}$ is the maximum leaf-specific whole-plant conductance and P(ψ) is a downregulating function of *k* that represents the phenomenological process of xylem embolism and outside-xylem decline in conductance as water potential decreases. P(ψ), also known as vulnerability curve, is represented here by a cumulative Weibull distribution

$$P(\psi)=\left(\frac{1}{2}\right)^{\left(\frac{\psi}{\psi_{50}}\right)^b} \qquad \text{(S4)}$$

where ψ is the water potential at a certain point within the water pathway, $\psi_{50}$ is the water potential at which a 50% of the conductance is lost, and *b* is the curve's shape. Notice that as defined in equation 2, Q is thus the integration of the conductance of each infinitesimal element of the water pathway from the soil to outside the xylem in the leaves assuming a common whole-plant vulnerability curve.

To satisfy water balance at steady-state, the supply function and the transpiration function must be equal (i.e. Q = E), then, $g_s$ and $\psi_l$ can be linked combining equations 1 and 2,

$$g_s=\frac{K_{max}}{1.6\, D\eta}\int_{\psi_s}^{\psi_l} P(\psi)\, d\psi \qquad \text{(S5)}$$

Photosynthesis model

Water transport and photosynthesis are directly linked by the rate of diffusion of $CO_2$ into the leaves. We can assume that at steady-state the rate of $CO_2$ entering the leaf is equal to the rate of $CO_2$ fixed by photosynthesis (A). This rate is also defined by the Fick's law as,

$$A = g_s (c_a - c_i) = g_s c_a (1 - \chi) \qquad \text{(S6)}$$

where $c_a$ and $c_i$ are the atmospheric and the leaf-internal partial pressures of $CO_2$, respectively, and $\chi$ is the ratio of $c_i$ over $c_a$.

Independently, leaf carbon assimilation rate in $C_3$ plants can be accurately described by the Farquhar-von Caemmerer-Berry model (Farquhar et al., 1980). The model mechanistically represents leaf photosynthesis as a process limited by either the rate of carboxylation of ribulose-1,5-biphosphate (RuBP) or by the rate of RuBP regeneration following the rate of the electron transport chain in a light-dependent reaction. The carboxylation-limited rate is defined as,

$$A_c = V_{cmax} \frac{c_i - \Gamma^*}{c_i + K_M} - b_0 V_{cmax} \qquad \text{(S7)}$$

where $V_{cmax}$ is the maximum carboxylation rate capacity, $\Gamma^*$ is the $CO_2$ compensation point without mitochondrial respiration, $K_M$ is the Michaelis-Menten constant for carboxylase activity of Rubisco in $C_3$ plants, and $b_0$ is the day-respiration rate per unit of $V_{cmax}$, as we assume day-respiration ($R_d$) proportional to carboxylation capacity (i.e., $R_d = b_0 V_{cmax}$) with $b_0 = 0.015$. Both $\Gamma^*$ and $K_M$ coefficients are a function of temperature ($T$) and atmospheric pressure ($p$) (Stocker et al., 2020). On the other side, the electron transport-limited rate is defined as,

$$A_j = \frac{J}{4} \left( \frac{c_i - \Gamma^*}{c_i + 2\Gamma^*} \right) - b_0 V_{cmax} \qquad \text{(S8)}$$

where $J$ is the effective electron-transport rate. Since $J$ saturates as light availability increase it can be approximated using a non-rectangular hyperbola,

$$J=\frac{4\phi_0 I_{abs}}{\sqrt{1+\left(\frac{4\phi_0 I_{abs}}{J_{max}}\right)^2}} \quad \text{(S9)}$$

here, $I_{abs}$ is the absorbed photosynthetic photon flux density (PPFD), $\phi_0$ is the intrinsic quantum yield, and $J_{max}$ is the maximum electron-transport rate achieved at $I_{abs} = \infty$.

Generally, the rate of carbon assimilation is assumed to be the minimum between those obtained by the two limiting processes, *i.e.* min($A_c$, $A_j$). In this study we assume the photosynthetic coordination hypothesis in order to compare the capacity of models with and without photosynthetic acclimation. It is assumed that since carboxylation and electron transport capacities are expensive to maintain, $A_c$ and $A_j$ are coordinated as a result of an acclimation process on a weekly to monthly timescales (Maire et al., 2012; Sabot et al., 2022a). This coordination implies that,

$$A_c=A_j=V_{cmax}\frac{c_i-\Gamma^*}{c_i+K_M}-b_0 V_{cmax}=\frac{J}{4}\left(\frac{c_i-\Gamma^*}{c_i+2\Gamma^*}\right)-b_0 V_{cmax} \quad \text{(S10)}$$

Then, $J$ can be expressed as a function of $V_{cmax}$ and $A_c$ as,

$$J=4V_{cmax}\frac{\left(c_i+2\Gamma^*\right)}{\left(c_i+K_M\right)}=\frac{4A_c\left(c_i-2\Gamma^*\right)}{c_i-\Gamma^*-b_0 V_{cmax}\left(c_i-2\Gamma^*\right)} \quad \text{(S11)}$$

and therefore obtain the coordinated $J_{max}$ by the inversion of equation 8,

$$J_{max}=\frac{4\phi_0 I_{abs}}{\sqrt{\left(\frac{4\phi_0 I_{abs}}{J}\right)^2-1}} \quad \text{(S12)}$$

Similarly, the coordinated $V_{cmax}$ can be obtained from $J_{max}$ or $A_j$,

$$V_{cmax}=\frac{J}{4}\frac{\left(c_i-K_M\right)}{\left(c_i-2\Gamma^*\right)}=A_j\frac{\left(c_i+K_M\right)}{\left(c_i\left(1-b_0\right)-\left(\Gamma^*+K_M b_0\right)\right)} \quad \text{(S13)}$$

In order to standardise the values of $V_{cmax}$ and $J_{max}$ we used the Arrhenius Eqs. S14 and S15 respectively,

$$V_{cmax,T1}=V_{cmax,T2}\,e^{\left(\frac{H_a}{R}\right)\left(\frac{1}{T2}-\frac{1}{T1}\right)} \quad \text{(S14)}$$

$$J_{max,T1} = J_{max,T2}\, e^{\left(\frac{H_{aj}}{R}\right)\left(\frac{1}{T2} - \frac{1}{T1}\right)} \quad \text{(S15)}$$

were T1 and T2 are the reference temperature and the actual temperature in K, respectively; $H_a$ is 65330 J $mol^{-1}$ that is the activation energy of $V_{cmax}$; $H_{aj}$ is 43900 J $mol^{-1}$ that is the activation energy of $J_{max}$; and R is the gas constant (8.314 J $mol^{-1}$ $K^{-1}$).

### Figures

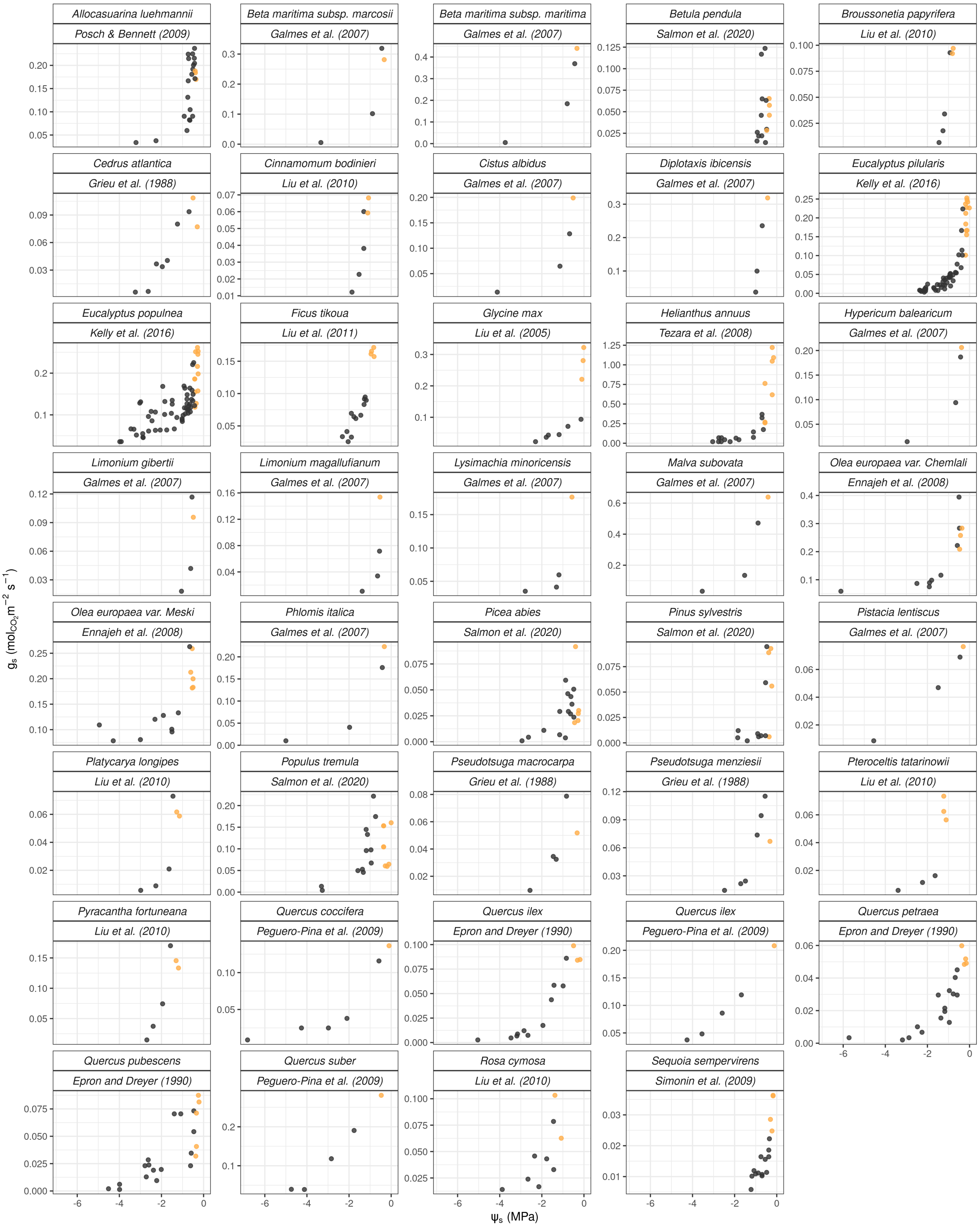


Figure S1. Scatter plots of actual $g_s$ from the species used to calibrate the models. Yellow dots present the species data of the $80^{th}$ wet percentile.

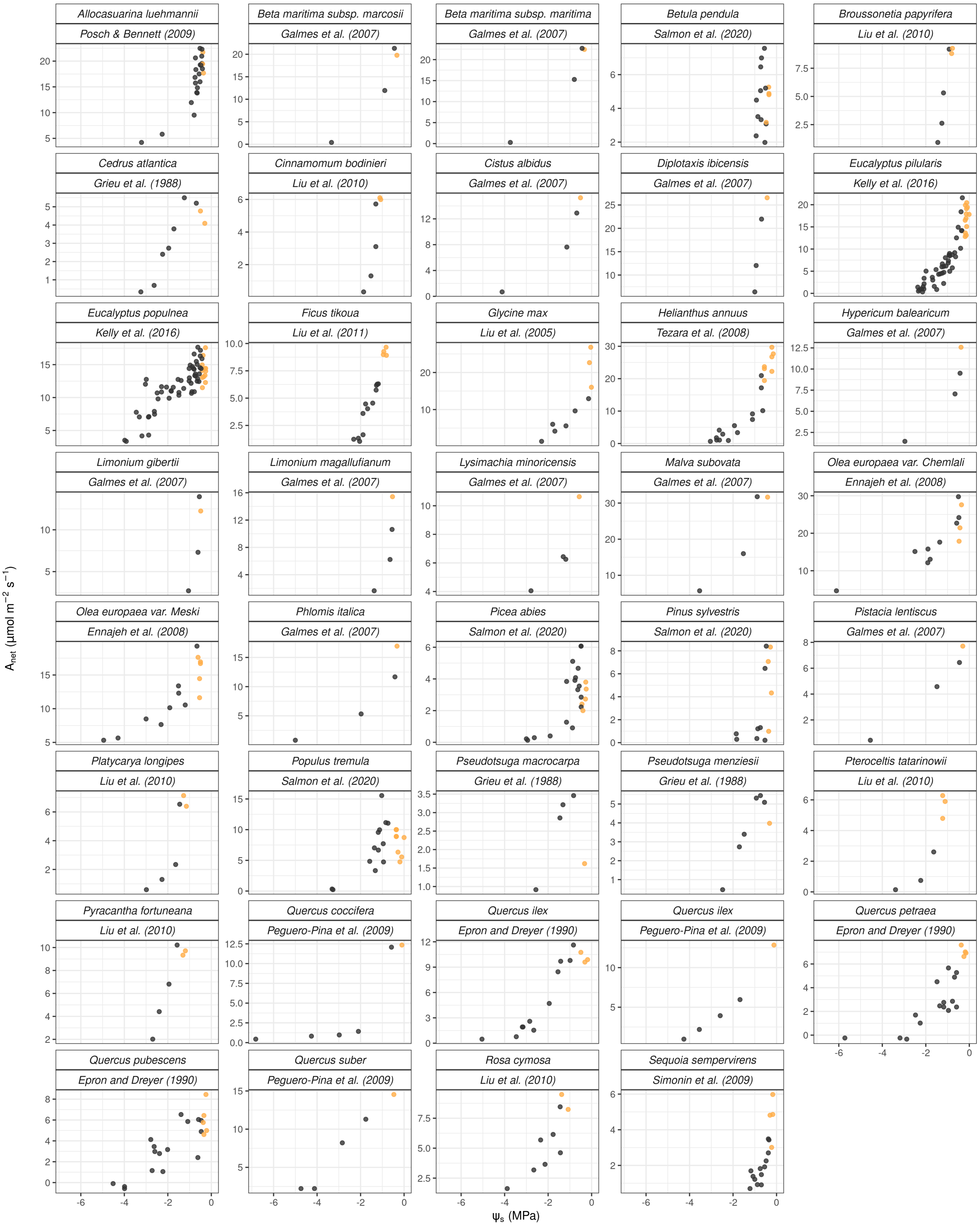


Figure S2. Scatter plots of actual $A_{net}$ from the species used to calibrate the models. Yellow dots present the species data of the 80$^{th}$ wet percentile.

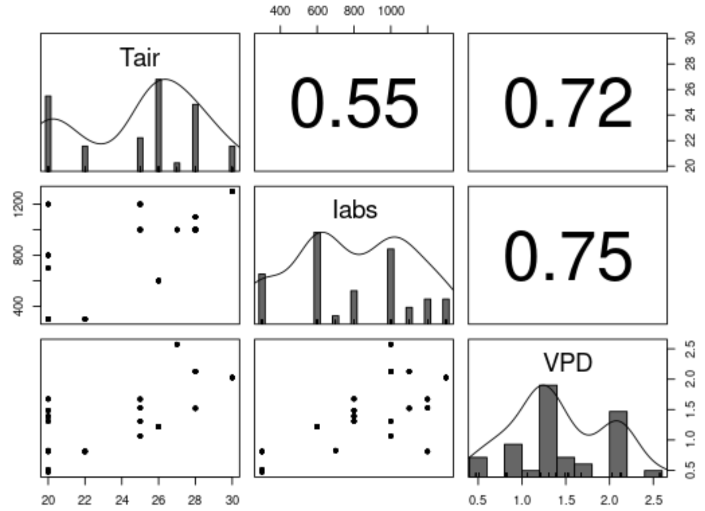


Figure S3. Bivariate scatter plots (lower diagonal) and Pearson's correlation (upper diagonal) pairs of average air temperature (Tair), absorbed photosynthetic active flux radiation (Iabs) and vapour pressure deficit (VPD) of the experimental periods. Data distributions are presented in the diagonal panels. Tair is in ºC, Iabs is in µmol $m^{-2}$ $s^{-1}$, and VPD is in kPa.

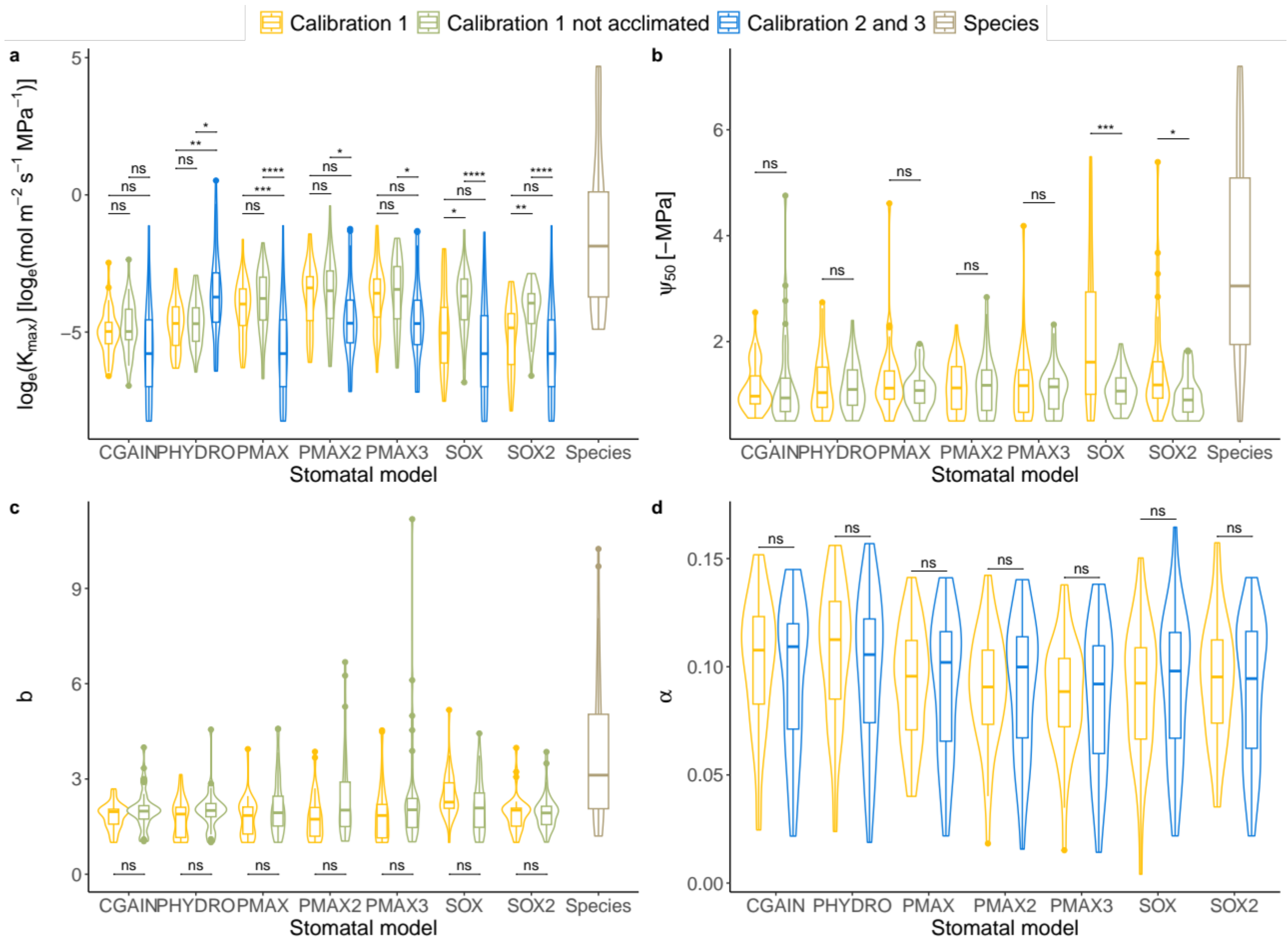


Figure S4. Distributions of the calibrated parameters $K_{max}$, $\psi_{50}$ , *b* and α and the extracted from the literature $K_{max}$, $\psi_{50}$ , *b* (Table 1). We present intra-model differences in parameter distributions across calibration approaches, that were estimated using a t-test model and adjusting p-values using Bonferroni. For calibration 1, all four parameters were calibrated, and an additional specific hydraulic parameter was calibrated for CGAIN, PHYDRO and SOX2. For calibration 1 not acclimated we calibrated $K_{max}$, $\psi_{50}$ , *b* and the corresponding specific parameter in CGAIN, PHYDRO and SOX2. In calibrations 2 we calibrated $K_{max}$ without acclimation and used the species specific $\psi_{50}$ , *b* extracted from the literature. In calibration 3, we used the $K_{max}$ calibrated in calibration 2 and calibrated α. $K_{max}$ units were transformed to mol $m^{-1}$ $s^{-1}$ $MPa^{-1}$. Statistical levels of significance: ns not significant; * p-value < 0.05; *** p-value < 0.01; *** p-value < 0.001; **** p-value < 0.0001.

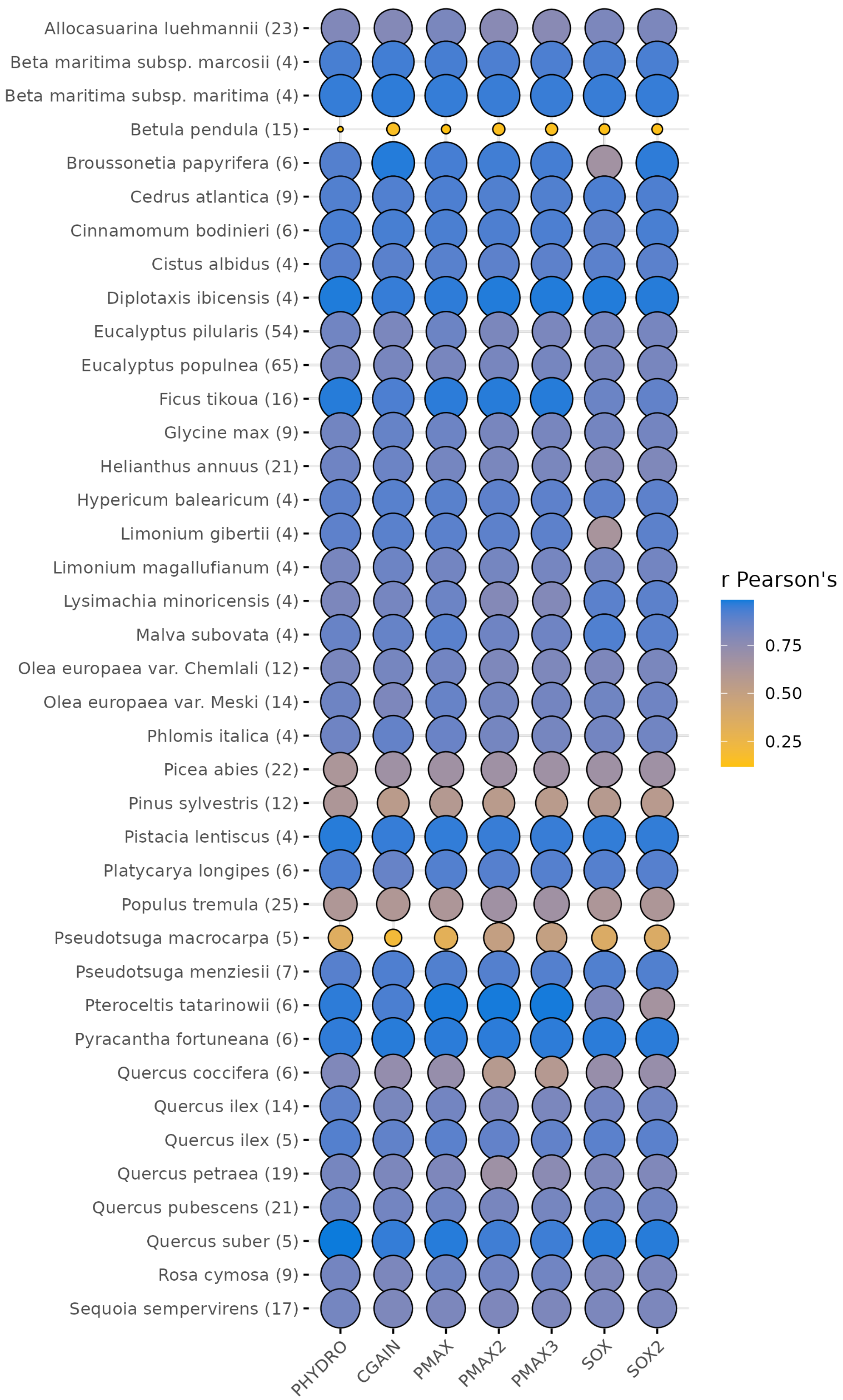


Figure S5. $A_{net}$ Pearson's Correlation for each species and stomatal model using Calibration 2 parameters with average α. N for each species is given in brackets.

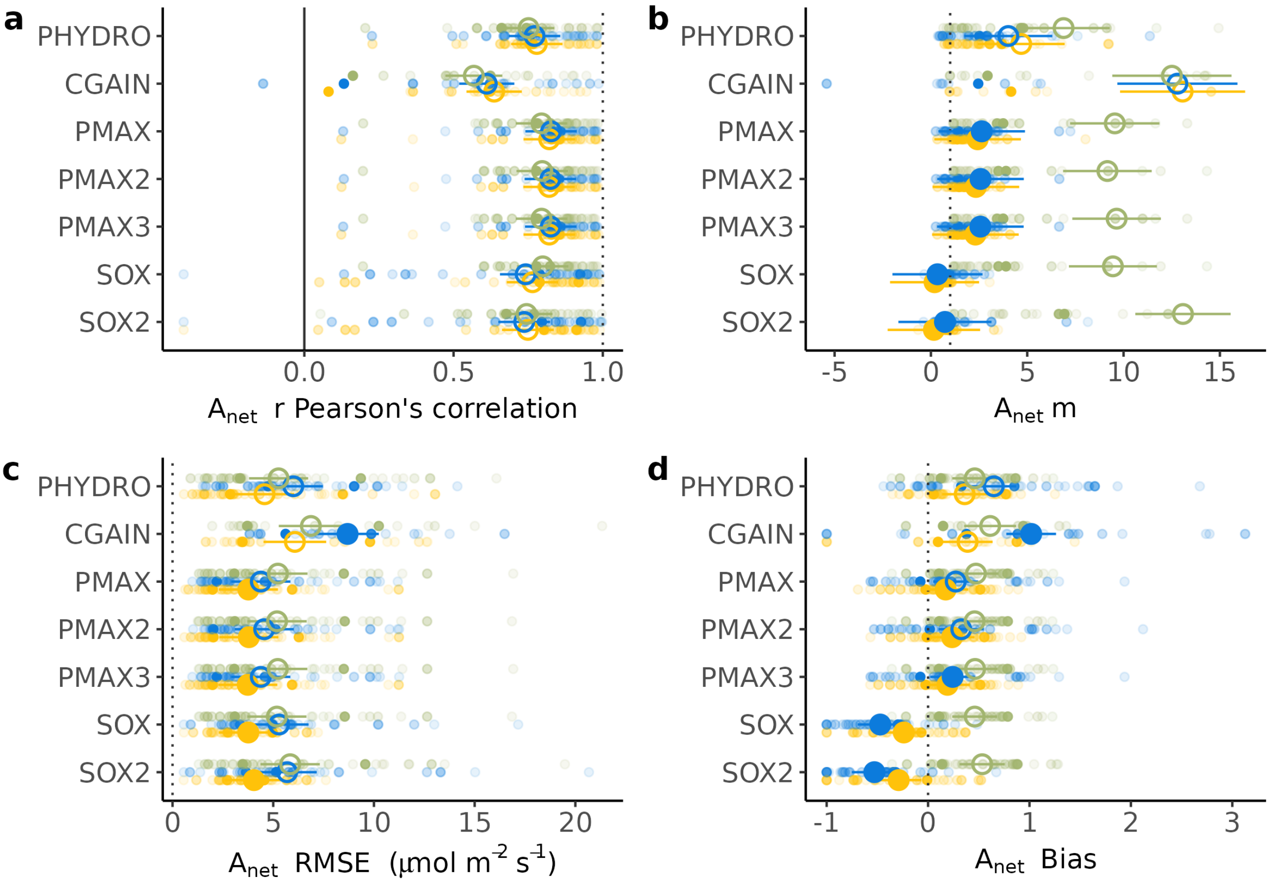


Figure S6. Performance comparison between acclimated and non-acclimated stomatal optimisation models, relative to species-specific observations of $A_{net}$ during different dry-down experiments using alternative vulnerability curve parameters. All the performance metrics were calculated for estimates of $A_{net}$ obtained within their full data sample, having fitted just one hydraulic parameter per stomatal model on the 80$^{th}$ wet percentile of the species data samples (not acclimated), then acclimating by setting α to 0.0945 for all the species (average α), and finally acclimating with a calibrated α. For PHYDRO, we used the species-level vulnerability curve parameters $\psi_{50}$ and *b* shown in Table 1. For all the other models, except PHYDRO, we used species-level outer xylem vulnerability curve parameters approximated as $\psi_{ox50} = \psi_{50}/3$ and $b = 1$. Small circles are the values for each species within each dry-down experiment, higher transparency indicates smaller data sample. Large circles are the metric averages estimated using an LMM (see Methods). Large closed circles indicate a significant difference in the metric averages of the model realisations with acclimation with respect to not acclimated ones; large open circles indicate no significant difference. Statistical significance was calculated using paired t-test comparisons. Vertical dotted lines indicate the best achievable performance for each metric. a) Pearson's correlation b) slope (*m*) of the linear relationship between the actual and estimated $g_s$ c) RMSE d) bias of the estimation of $g_s$ calculated as in Eq. 10.

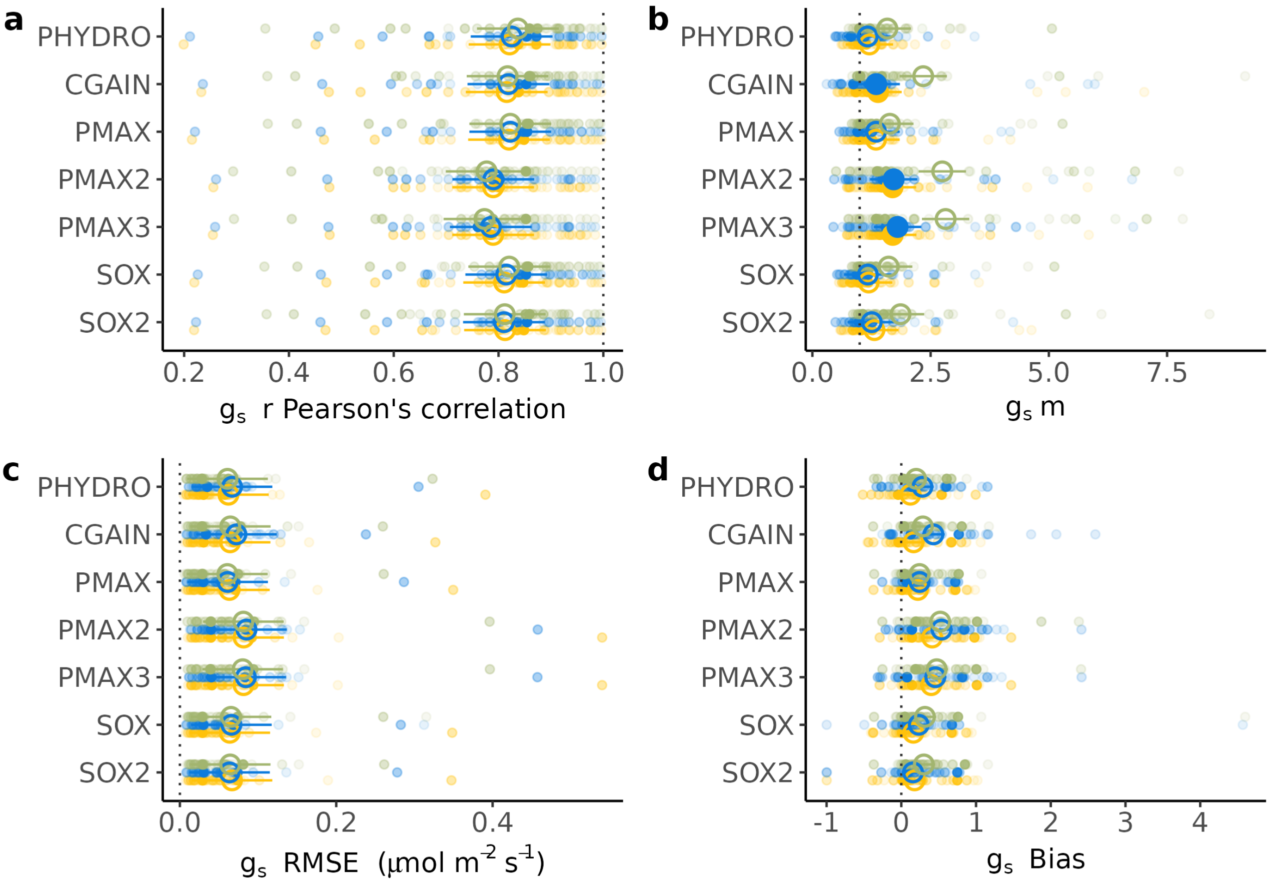


Figure S7. Performance comparison between acclimated and non-acclimated stomatal optimisation models, relative to species-specific observations of $g_s$ during different dry-down experiments. All the performance metrics were calculated for estimates of $g_s$ obtained within their full data sample, having fitted just one hydraulic parameter per stomatal model on the $80^{th}$ wet percentile of the species data samples (not acclimated), then acclimating by setting α to 0.0945 for all the species (average α), and finally acclimating with a calibrated α. For all the models, except PHYDRO, we used the species-level vulnerability curve parameters $\psi_{50}$ and *b* shown in Table 1. For PHYDRO we used species-level outer xylem vulnerability curve parameters as $\psi_{ox50} = \psi_{50}/3$ and $b = 1$. Small circles are the values for each species within each dry-down experiment, higher transparency indicates smaller data sample. Large circles are the metric averages estimated using an LMM (see Methods). Large closed circles indicate a significant difference in the metric averages of the model realisations with acclimation with respect to not acclimated ones; large open circles indicate no significant difference. Statistical significance was calculated using paired t-test comparisons. Vertical dotted lines indicate the best achievable performance for each metric. a) Pearson's correlation b) slope (*m*) of the linear relationship between the actual and estimated $g_s$ c) RMSE d) bias of the estimation of $g_s$ calculated as in Eq. 10.

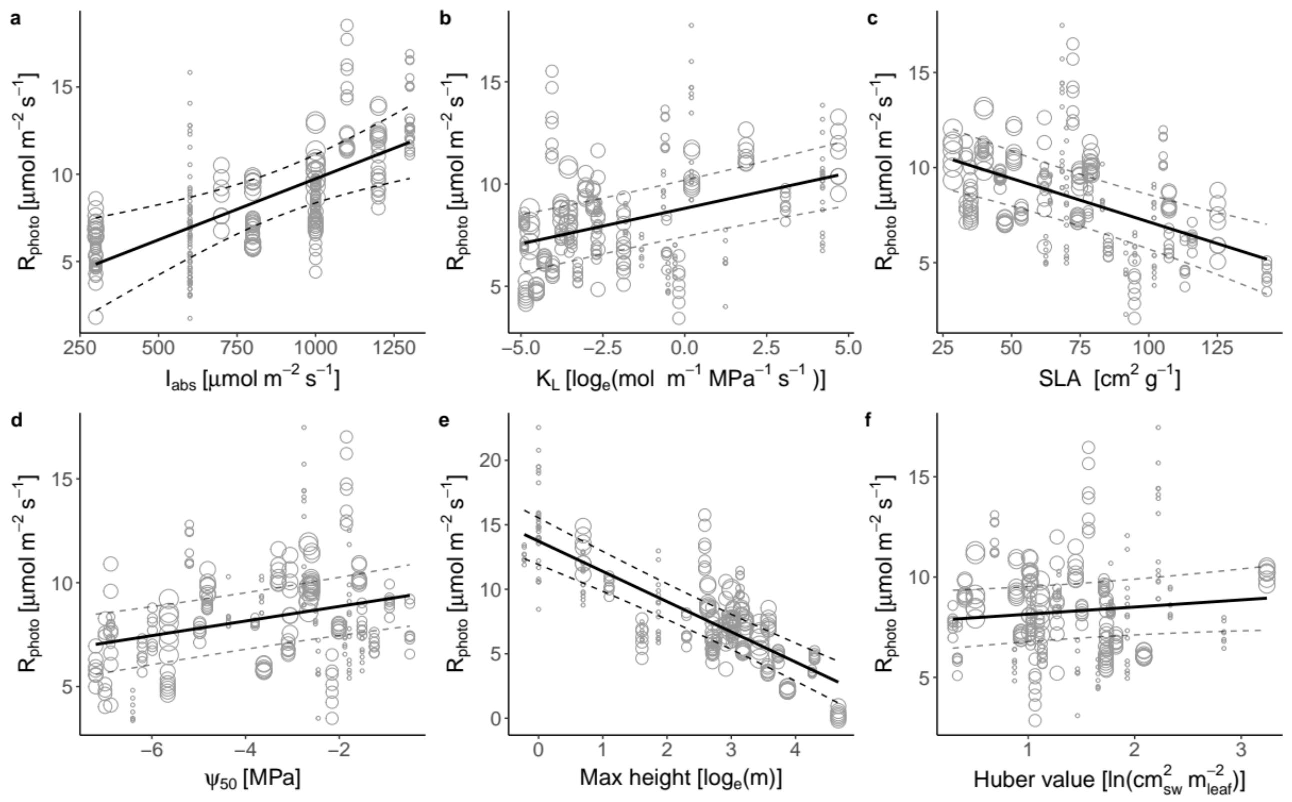


Figure S8. Partial effects from a multivariate Linear-Mixed Model (LMM) explaining the variability of photosynthetic capacity cost under well-watered conditions (or photosynthetic maintenance respiration; $R_{photo} = \alpha J_{maxWW,25}$). The LMM was fitted using the dry-down experiments and stomatal models as crossed random intercepts. Grey scatter points show all the model-specific contributions to α, across all species. Black plain lines are partial regressions and dashed lines represent the partial regressions plus or minus their standard error. Size of the points represent the weight applied to each data point, which is equal to the natural logarithm of the sample size.

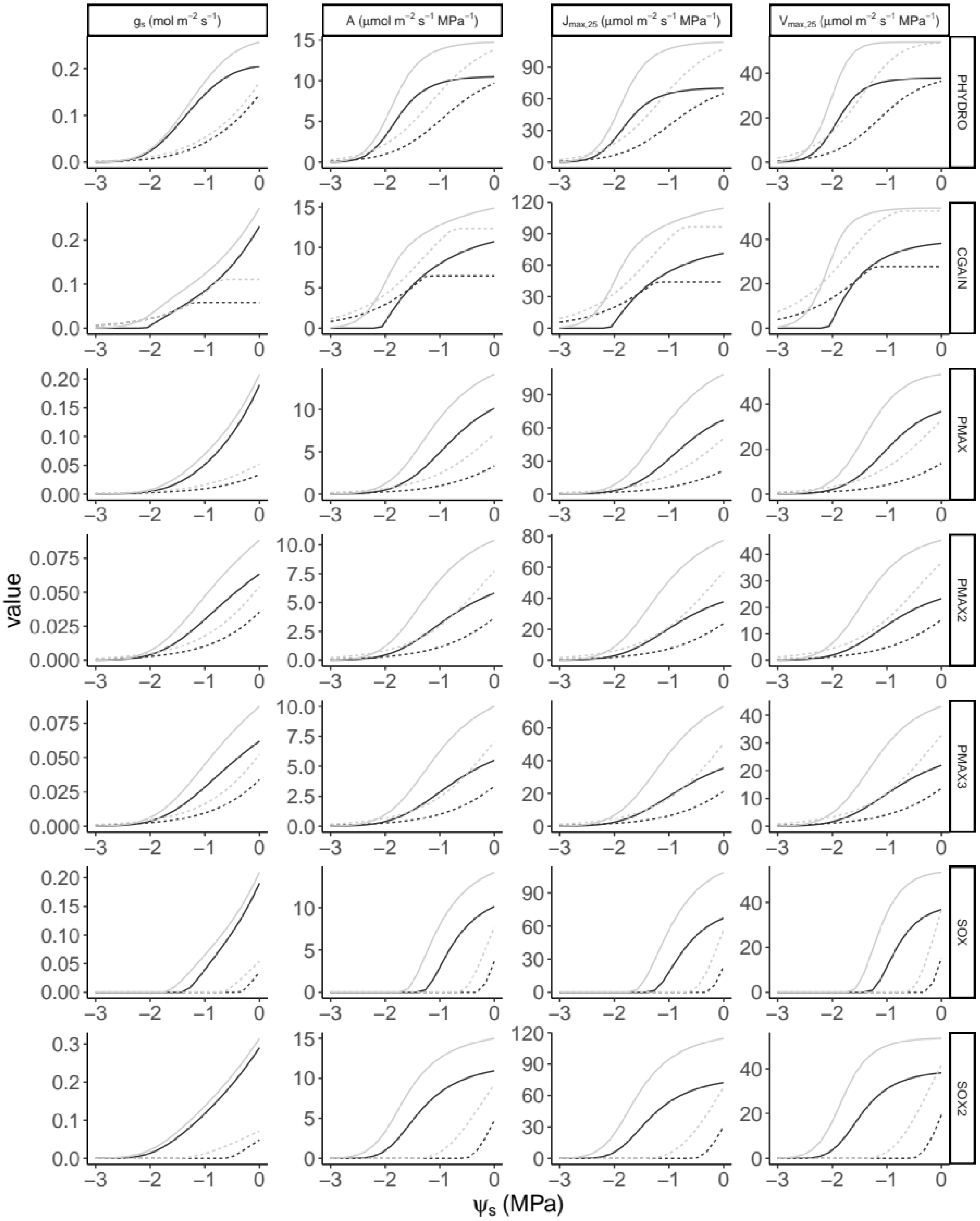


Figure S9. Example of the shape of each acclimated stomatal optimization model using equivalent parameters and different levels of α and $\psi_{50}$. Environmental variables with the exception of $\psi_s$ were maintained constant. Black lines present models with α = 0.0745 while grey lines are models with α = 0.1145. We use *Ficus tikoua* vulnerability curve parameters as reference. Solid lines are obtained with $\psi_{50}$ = -1.58 MPa and dashed lines with $\psi_{ox50}$ = $\psi_{50}/3$ MPa and $b$ = 3.12. For all the models and simulations $K_{max}$ = 0.0062 $mol_{H2O}$ $m_{leaf}^{-2}$ $s^{-1}$ $MPa^{-1}$ . PHYDRO parameter $\gamma$ = 1 µmol $m_{leaf}^{-2}$ $s^{-1}$ $MPa^{-2}$, SOX2 parameter $\delta$ = 2, and CGAIN parameter $\varpi$ = 5 µmol $m_{leaf}^{-2}$ $s^{-1}$. The environmental variables used were set to: $T_a$ = 25 ºC, PPFD = 600 µmol $m_{leaf}^{-2}$ $s^{-1}$, VPD = 1000 Pa, $C_a$ = 400 ppm, FAPAR = 0.99, quantum yield efficiency parameter $\varphi_0$ = 0.087.

**Tables**

Table S1. RMSE of traits obtained using imputation methodology. The total number of rows (N), number of rows with observed data (N Obs), number of rows with gaps (N Gaps), and Root Mean Square Error (RMSE) of the gap-filling process are presented. $H_{max}$ is the maximum height of the species in m. The variable Hv is the Huber value or sapwood area per leaf area in $cm_{sw}^2$ $m_{leaf}^{-2}$. $\psi_{12}$, $\psi_{50}$, and $\psi_{88}$ are the water potentials at 12%, 50%, and 88% loss of xylem hydraulic conductivity, respectively, at the species level in MPa. $K_L$ is the maximum leaf-area specific hydraulic conductivity in mol $m_{leaf}^{-1}$ $s^{-1}$ $MPa^{-1}$. SLA is the specific leaf area at the species level in $cm^2$ $g^{-1}$.

| Variable | N | N Obs | N Gaps | RMSE |
|---|---|---|---|---|
| $H_{max}$ | 1635 | 813 | 822 | 9.523 |
| Hv | 1635 | 434 | 1201 | 0.001 |
| $K_L$ | 1635 | 559 | 1076 | 0.466 |
| $\psi_{12}$ | 1635 | 922 | 713 | 1.393 |
| $\psi_{50}$ | 1635 | 1220 | 415 | 1.564 |
| $\psi_{88}$ | 1635 | 957 | 678 | 2.063 |
| SLA | 1635 | 378 | 1257 | 36.154 |

Table S2. Estimated marginal means of $A_{net}$ RMSE metric ($\mu mol_C$ $m^{-2}$ $s^{-1}$) from not acclimated, the average α acclimated (calibration 2) and the calibrated α acclimated (calibration 3) stomatal models. Confident level (CL) and P-value adjustment using Sidak method with the R package emmeans (Lenth, 2022; Searle et al., 1980). Different letters imply significant differences between groups ($p<0.05$) within each calibration type. Stomatal models are ordered by calibration and from lower to higher RMSE

| Stomatal model | Calibration type | RMSE | Lower CL | Upper CL | Group |
|---|---|---|---|---|---|
| PHYDRO | Not acclimated | 4.69 | 2.40 | 6.97 | a |
| CGAIN | Not acclimated | 5.21 | 2.92 | 7.49 | a |
| PMAX | Not acclimated | 4.93 | 2.64 | 7.21 | a |
| PMAX2 | Not acclimated | 5.49 | 3.21 | 7.78 | a |

| | | | | | |
|---|---|---|---|---|---|
| PMAX3 | Not acclimated | 5.45 | 3.17 | 7.74 | a |
| SOX | Not acclimated | 4.94 | 2.66 | 7.23 | a |
| SOX2 | Not acclimated | 5.18 | 2.90 | 7.47 | a |
| PHYDRO | Average α | 5.17 | 2.89 | 7.46 | ab |
| CGAIN | Average α | 5.61 | 3.33 | 7.90 | b |
| PMAX | Average α | 4.24 | 1.95 | 6.52 | a |
| PMAX2 | Average α | 5.01 | 2.73 | 7.29 | ab |
| PMAX3 | Average α | 4.79 | 2.50 | 7.07 | ab |
| SOX | Average α | 4.19 | 1.90 | 6.47 | a |
| SOX2 | Average α | 4.38 | 2.09 | 6.66 | a |
| PHYDRO | Calibrated α | 3.63 | 1.35 | 5.92 | a |
| CGAIN | Calibrated α | 4.10 | 1.81 | 6.38 | a |
| PMAX | Calibrated α | 3.74 | 1.45 | 6.02 | a |
| PMAX2 | Calibrated α | 4.32 | 2.03 | 6.60 | a |
| PMAX3 | Calibrated α | 4.26 | 1.98 | 6.55 | a |
| SOX | Calibrated α | 3.80 | 1.52 | 6.09 | a |
| SOX2 | Calibrated α | 3.88 | 1.59 | 6.16 | a |

Table S3. Estimated marginal Coefficient of Determination ($R^2$) of gs and $A_{net}$ from acclimated and non-acclimated stomatal models across species using the best set of parameters. Calculated using a LMM with the actual $g_s$ or $A_{net}$ value as predicted variable and the estimated $g_s$ or $A_{net}$ as fixed explanatory variable. We included in the LMM's both random slopes of $g_s$ or $A_{net}$, and random intercepts for each species.

| | Not acclimated | | Acclimated | |
|---|---|---|---|---|
| Stomatal model | $R^2$ $g_s$ | $R^2$ $A_{net}$ | $R^2$ $g_s$ | $R^2$ $A_{net}$ |
| PHYDRO | 0.728 | 0.642 | 0.725 | 0.704 |
| CGAIN | 0.739 | 0.644 | 0.727 | 0.721 |
| PMAX | 0.717 | 0.621 | 0.722 | 0.712 |
| PMAX2 | 0.535 | 0.620 | 0.583 | 0.702 |
| PMAX3 | 0.527 | 0.625 | 0.634 | 0.713 |
| SOX | 0.719 | 0.622 | 0.716 | 0.720 |
| SOX2 | 0.708 | 0.639 | 0.576 | 0.689 |
